# Lipogenesis and innate immunity in hepatocellular carcinoma cells reprogrammed by an isoenzyme switch of hexokinases

**DOI:** 10.1101/2020.03.13.973321

**Authors:** Laure Perrin-Cocon, Pierre-Olivier Vidalain, Clémence Jacquemin, Anne Aublin-Gex, Keedrian Olmstead, Baptiste Panthu, Gilles J. P. Rautureau, Patrice André, Piotr Nyczka, Marc-Thorsten Hütt, Nivea Amoedo, Rodrigue Rossignol, Fabian Volker Filipp, Vincent Lotteau, Olivier Diaz

**Author notes:** These authors contributed equally to this work (Laure Perrin-Cocon, Pierre-Olivier Vidalain). These authors jointly supervised this work (Vincent Lotteau, Olivier Diaz).

## Abstract

During the cancerous transformation of normal hepatocytes into hepatocellular carcinoma (HCC), the enzyme catalyzing the first rate-limiting step of glycolysis, namely the glucokinase (GCK), is replaced by the higher affinity isoenzyme, hexokinase 2 (HK2). The transcriptomic analysis of HCC tumors shows that highest expression level of *HK2* in tumor lesions is inversely correlated to *GCK* expression, and is associated to poor prognosis for patient survival. To further explore functional consequences of the GCK-to-HK2 isoenzyme switch occurring during carcinogenesis, *HK2* was knocked-out in the HCC cell line Huh7 and replaced by *GCK*, to generate the Huh7-*GCK*^*+*^/*HK2*^*−*^ cell line. HK2 knockdown and GCK expression rewired central carbon metabolism, stimulated mitochondrial respiration and restored essential metabolic functions of normal hepatocytes such as lipogenesis, VLDL secretion, glycogen storage. It also reactivated innate immune responses and sensitivity to natural killer cells, showing that consequences of the HK switch extend beyond metabolic reprogramming.

## Introduction

Hepatocellular carcinoma (HCC) is the most common liver cancer and the fourth leading cause of cancer-related death (1). HCC is closely linked to chronic liver inflammation, chronic viral hepatitis, exposure to toxins, and metabolic dysfunction such as non-alcoholic steatohepatitis (NASH). HCC is of poor prognosis, and treatments are essentially based on surgical resection, liver transplantation or aggressive chemo and/or radiotherapy. In patients with advanced HCC, broad-spectrum kinase inhibitors are approved (2) but with limited benefit (3). Effective personalized therapies are needed but their development is impeded by our poor understanding of molecular mechanisms underlying HCC onset and progression. Efforts to characterize the disease on the basis of etiology and outcomes revealed metabolic deregulation as a hallmark of HCC progression (4). Indeed, metabolic remodeling is critically required for tumor growth, since bioenergetic requirements and anabolic demands drastically increase (5–7). For instance, HCC cells have lost their ability to secrete very low-density lipoproteins (VLDL), a highly specialized function of hepatocyte and can only secrete low-density lipoproteins (LDL)-like lipoproteins, indicating a defective lipogenesis and/or lipoprotein assembly (8).

Metabolic reprogramming in cancer cells involves the modulation of several enzymes by oncogenic drivers (6). Targeting these enzymes is now considered as a therapeutic strategy for several types of cancers (6). Among these enzymes, hexokinase 2 (HK2) stands out because of its elevated or induced expression in numerous cancers, including HCC (9). Hexokinases control the first rate-limiting step of glucose catabolism by phosphorylating glucose to glucose-6-phosphate (G6P), fueling glycolysis as well as glycogen, pentose phosphate and triglyceride synthesis. The human genome contains four genes encoding distinct hexokinase isoenzymes, named HK1 to HK4 (HK4 is also known as glucokinase or GCK), with distinct enzymatic kinetics and tissue distributions. A fifth putative hexokinase enzyme was recently discovered but has not been fully characterized yet (10). A switch from GCK to HK2 isoenzymes is occurring during the transition from primary to tumor hepatocytes so that HCC cell lines express HK2 but no longer GCK. HK2 expression level has been correlated with disease progression and dedifferentiation of HCC cells (11). When HK2 is artificially knocked-down in HCC cell lines, glycolysis is repressed, and tumorigenesis is inhibited while cell death increases (9). In addition, hexokinase function extends beyond metabolism towards autophagy, cell migration, and immunity, suggesting that the GCK-to-HK2 isoenzyme switch has broader consequences than initially suspected (12–15). Here, we analyzed transcriptomic data of HCC biopsies and correlated hexokinase isoenzyme expression level with patient survival. This led us to generate a new human HCC model expressing *GCK* instead of *HK2*. A comparative analysis of *GCK*^*+*^ vs *HK2*^*+*^ HCC cell lines provided a unique opportunity to look into HK isoenzyme-dependent metabolic features, lipoprotein production and resistance to immune signals of liver cancer cells.

## Results

### Relative expression level of GCK and HK2 in HCC patients

Although an isoenzyme switch from GCK to HK2 has been observed during the carcinogenesis process (16), whether hexokinase isoenzymes expression is predictive of patient survival is unclear. We first analyzed the transcriptomes (RNAseq data) of 365 HCC biopsies from The Cancer Genome Atlas (TCGA) database (17, 18)(Supplementary Table 1). For each HK, the individual gene expression level was used to stratify patients into two subgroups according to Uhlen et al. (18) and overall survival in the two subgroups was determined using a Kaplan-Meier’s estimator. Although *HK1* or *HK3* expression level were not associated to patient survival rate (Fig. 1a), highest expression levels of *HK2* as previously described (19) and lowest expression levels of *GCK* in the tumors were associated with a lower survival rate. We thus stratified patients based on the GCK/HK2 expression ratio to combine these two markers (Fig. 1b). When patients were stratified on the basis of *HK2* or *GCK* expression levels, the median survival between the corresponding subgroups differed by 33.8 and 36.5 months, respectively (Fig. 1a). This difference reached 42.8 months when the stratification of patients was based on the *GCK/HK2* ratio (Fig. 1b). This demonstrated that the *GCK/HK2* ratio outperforms *HK2* or *GCK* expression alone as predictor of patient survival. Finally, correlation coefficients between patient survival in months and *HK2* or *GCK* expression level were determined. For this, we only considered the subset of 130 patients for whom the period between diagnosis and death is precisely known (uncensored data), and performed a Spearman’s rank correlation test (Fig. 1c). Patient survival was positively correlated to *GCK* expression but inversely correlated to *HK2* expression in line with the Kaplan-Meier analysis. In addition, *GCK* and *HK2* expression tends to be inversely correlated in tumor samples (Fig. 1c). Therefore, there is a trend for mutual exclusion of *GCK* and *HK2* expression in HCC tumors, and this profile is associated to clinical outcome.

**Figure 1.**
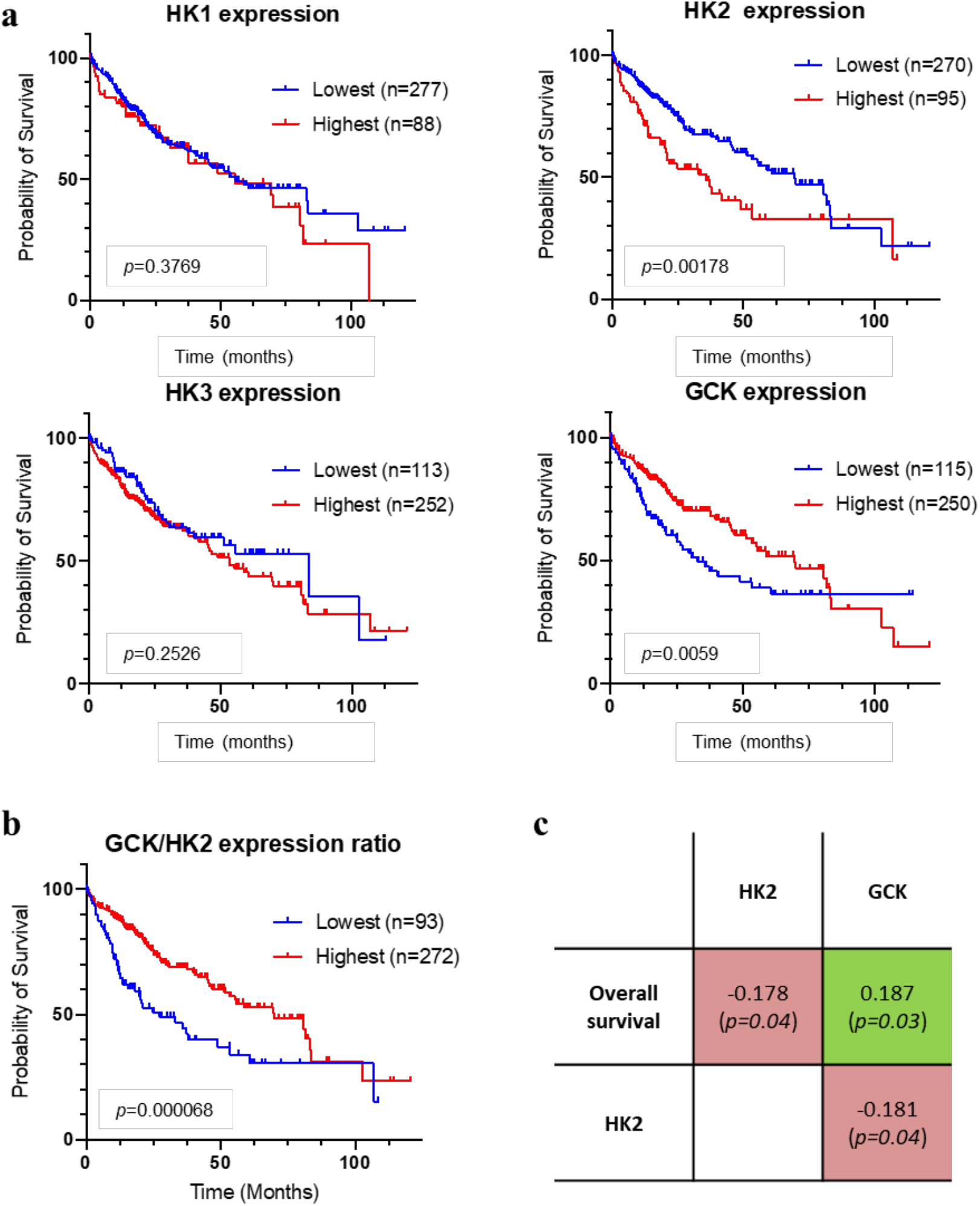
Correlation between hexokinase expression levels in HCC tumors and patient survival. **a** Kaplan–Meier estimates of the survival of HCC patients depending on the expression of HK1, HK2, HK3 and GCK (HK4) genes in tumor biopsies (n=365; TCGA expression data retrieved from cBioPortal)(70, 71). Duplicate analyses from the same patient were removed as well as patients who died when biopsied (overall survival=0 months). Optimal stratification based on highest and lowest gene expression values was determined using Protein Atlas database (18). **b** Same as above but patients were stratified based on the GCK/HK2 gene expression ratio. The stratification showing the lowest *p* value when comparing subgroups of patients with the highest to the lowest GCK/HK2 expression ratio is displayed. Patient TCGA-DD-AAE9 exhibiting undetectable levels of GCK and HK2 was removed from this analysis as the GCK/HK2 ratio could not be calculated. **c** Correlations between patient survival, GCK expression and HK2 expression. Spearman’s rank correlation test on the subset 130 patients for whom the period between diagnosis and death is precisely known (uncensored data).

### Engineering a cellular model of the hexokinase isoenzyme switch

To decipher functional consequences of GCK or HK2 expression in a HCC model, we restored GCK expression by lentiviral transduction in the reference HCC cell line Huh7, and knocked-out the endogenous *HK2* gene by CRISPR/Cas9. The exclusive expression of HK2 and *GCK* in Huh7 and Huh7-*GCK*^*+*^/*HK2*^*−*^ cell lines, respectively, was validated, while HK1 and HK3 were not expressed (Fig. 2a). The hexokinase activity in the presence of increasing concentration of glucose was determined in protein lysates from the two respective cell lines. Hexokinase activity in Huh7 lysate reached its maximum at low glucose concentration, presenting a saturation curve according to Michaelis–Menten kinetics (Fig. 2b). In contrast, the hexokinase activity in Huh7-*GCK*^*+*^/*HK2*^*−*^ lysates followed a pseudo-allosteric response to glucose (20, 21). Thus, the expected HK2 and GCK activities were observed in the Huh7 and Huh7-*GCK*^*+*^/*HK2*^*−*^ cells respectively. The cell proliferation capacity remained identical between the two cell lines (Supplementary Fig. 1). We then compared the genome edited Huh7-*GCK*^*+*^/*HK2*^*−*^ and the parental Huh7 cell lines (*i.e.* Huh7-*GCK*^*−*^/*HK2*^*+*^) at a transcriptomic, metabolic and immunological level.

**Figure 2.**
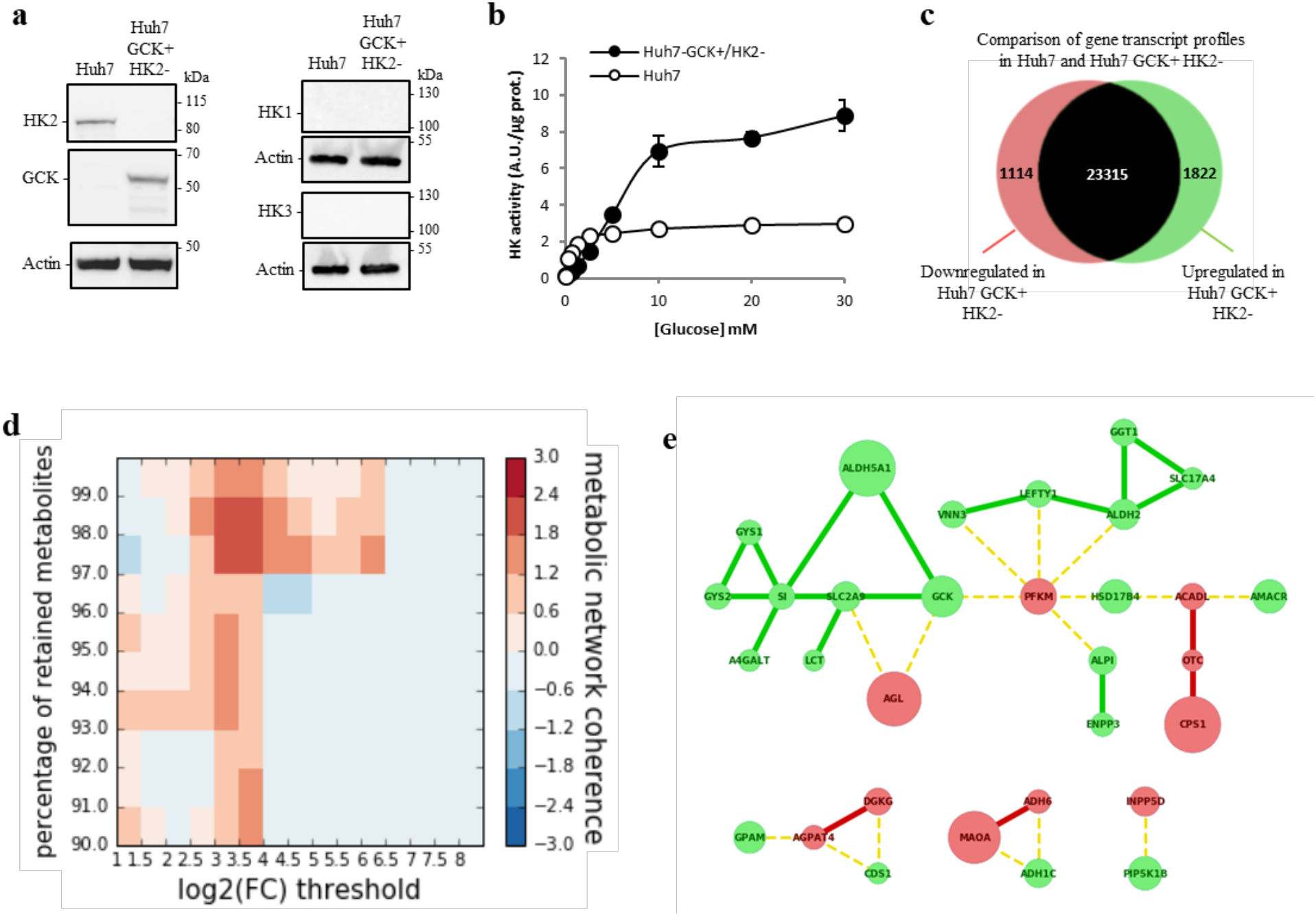
Hexokinase isoenzyme switch in Huh7 cells induces extended modifications of metabolic connections. **a** Western-blot analysis of HK1, HK2, HK3 and GCK expression in Huh7 and Huh7-*GCK*^*+*^/*HK2*^*−*^. **b** Hexokinase activity in homogenates of Huh7 and Huh7-*GCK*^*+*^/*HK2*^*−*^ cells. **c** Number of genes changing their expression pattern in Huh7 and Huh7-*GCK*^*+*^/*HK2*^*−*^ cells (see Supplementary Table 2 for details). **d** Heatmap showing clustering enrichment scores of the networks obtained when mapping differentially expressed genes to the human metabolic model Recon2. Clustering enrichment scores from the highest in red to the lowest in blue were calculated for different gene expression thresholds (Log_2_|FC|) and percentages of retained currency metabolites. **e** Gene network corresponding to the maximal clustering enrichment score (Log_2_|FC|>3; removed currency metabolites = 2%). The transcription of nodes in green was upregulated and those in red downregulated in Huh7-*GCK*^*+*^/*HK2*^*−*^ compared to Huh7 cells. Plain edges mark coregulation between nodes and broken edges inverse regulation at the transcriptional level.

### Transcriptomic data revealed extended modifications of metabolic connections in Huh7-GCK+/HK2-

Transcriptomic profiles of Huh7 and Huh7-*GCK*^*+*^/*HK2*^*−*^ cells were determined by next generation sequencing (Supplementary Table 2). Overall, 4.2% of the gene transcripts were reduced and 6.9% were induced in Huh7-*GCK*^*+*^/*HK2*^*−*^ compared to Huh7 (Fig. 2c; |fold-change (FC)|>2 and *p* value<0.05). We first determined the metabolic consequences of the HK isoenzyme switch by mapping the differentially expressed genes onto the a well-established bipartite metabolic network Recon2, connecting gene products and metabolites (Supplementary Fig. 2-4) (22, 23). After trimming highest-degree metabolites as currency metabolites, clusters of genes that are both differentially expressed and connected by common metabolites emerged. Interestingly, we found that across a wide range of analysis parameters, including varying rates of currency metabolites and gene expression fold-change, the differentially expressed metabolic genes are substantially better connected than expected by chance (Fig. 2d). This highlights the specificity of the transcriptomic changes with respect to metabolic pathways. The spanned network presented in Figure 2e corresponds to a stringent fold-change threshold for transcriptomic data (log_2_(|FC|)>3) while removing 2 percent of highest-degree currency metabolites. This network shows connected components within glycolysis, but also across distant modules including the gamma-aminobutyric acid (GABA) shunt (ALDH5A1), urea cycle (CPS1, OTC), glycogen metabolism (GYS1, GYS2, AGL) and lipid synthesis (GPAM, AGPAT4, DGKG, CDS1, A4GALT) or degradation (ACADL, HSD17B4, AMACR). This analysis highlights the global impact of the HK isoenzyme switch that spreads beyond glycolysis across distant connected metabolic modules.

Enrichment of molecular and cellular functions in differentially expressed genes was also analyzed using Ingenuity Pathway Analysis (IPA). This revealed that cellular movement and lipid metabolism were the most affected functions (Fig. 3a and Supplementary Table 3). A closer look at these annotations pointed to differences in the migratory capacities (Fig. 3b) as well as lipid concentration and synthesis (Fig. 3c). The migratory capacities of Huh7 and Huh7-*GCK*^*+*^/*HK2*^*−*^ were compared using transwell-migration cell assays (Fig. 3d-e). Results showed a higher migratory capacity of Huh7-*GCK*^*+*^/*HK2*^*−*^ cells, in line with Kishore M. et al showing that GCK expression induced by pro-migratory signals controls the trafficking of CD4^+^/CD25^+^/FOXP3^+^ regulatory T (T_reg_) cells (15). To validate the differences in lipid metabolism highlighted by the transcriptomic analysis, intracellular content in neutral lipids was first determined with lipophilic dyes. As assessed by Oil-Red-O or BODIPY staining, an accumulation of intracellular neutral lipids was observed in Huh7-*GCK*^*+*^/*HK2*^*−*^ in comparison to Huh7 (Fig. 3f and 3g). The accumulation of neutral lipids in Huh7 expressing both *HK2* and *GCK* indicates that the phenotype is driven by *GCK* expression and not by *HK2* knockdown (Supplementary Fig. 5). Lipid accumulation upon GCK expression was also observed in Huh6 hepatoblastoma cells but not in epithelial kidney Vero cells, indicating that this phenomenon occurs in metabolically relevant cells (Supplementary Fig. 5).

**Figure 3.**
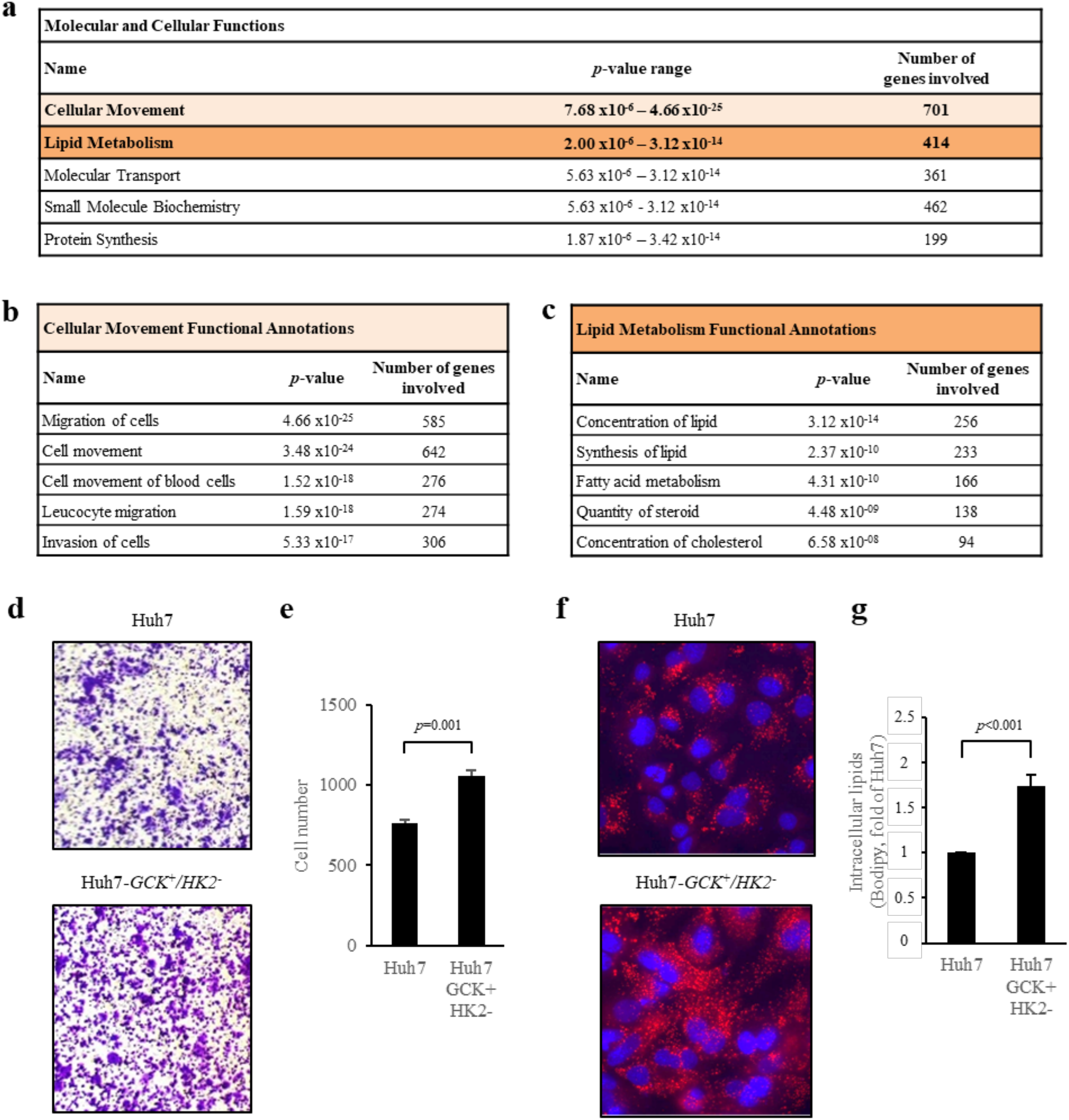
Huh7-*GCK*^*+*^/*HK2*^*−*^ cells have a higher migration capacity and lipid droplets content. **a** Analysis of differentially expressed genes in the two cell lines using gene set enrichment analysis (|FC|>2 with a *p* value <0.05). Top five enriched molecular and cellular functions are presented. **b** and **c** Ranked IPA-annotations associated with ‘cellular movement’ and ‘lipid metabolism’. **d, e** Results of transwell-migration tests. **d** Representative images and **e** count of migrating cells. **f** Oil Red-O staining of lipid droplets (red) with nucleus counterstaining (blue). **g** Quantification of intracellular lipids by FACS after BODIPY staining. E and G correspond to means ± SEM with n≥3 (*p* value determined by Student’s t-test).

### Differential lipid metabolism in Huh7 and Huh7-*GCK*^+^/*HK2*^−^

The intracellular lipid content of the two cell lines was further analyzed. In Huh7-*GCK*^*+*^/*HK2*^*−*^, an enrichment in phosphatidylcholine, cholesterol, triglycerides (TG) and free fatty acids was observed compared to Huh7 (Fig. 4a). One major function of hepatocytes is to secrete triglyceride-rich VLDL and this function is altered in HCC cells that secrete smaller lipoproteins with the density of LDL (24, 25). The secretion of lipids and lipoproteins by both cell lines was analyzed after a 24h-culture in the absence of FCS to exclude any participation of exogenous lipids in the production of lipoproteins. Huh7-*GCK*^*+*^/*HK2*^*−*^ secreted more free fatty acids than Huh7 while secretion of cholesterol and triglycerides (TG) remained unchanged (Fig. 4b). However, under the same conditions, the secretion of apolipoprotein B (ApoB) by Huh7-*GCK*^*+*^/*HK2*^*−*^ was reduced compared to Huh7. Since ApoB is a non-exchangeable protein with only one copy in VLDL and LDL particles, an elevated TG/ApoB ratio indicates that ApoB^+^-lipoproteins secreted by Huh7-*GCK*^*+*^/*HK2*^*−*^ cells are enriched in TG compared to those secreted by Huh7 (Fig. 4c). This was confirmed by the ApoB distribution in density gradient fractions. As expected, lipoproteins secreted by Huh7 sediment at the density of LDL, while those secreted by Huh7-*GCK*^*+*^/*HK2*^*−*^ (Fig. 4d) match the density of VLDL found in human plasma or secreted by primary human hepatocytes in culture (26, 27). This indicates that GCK expression is essential for the VLDL assembly/secretion pathway and could explain the loss of this crucial metabolic pathway in hepatoma cells expressing HK2 instead of GCK (28).

**Figure 4.**
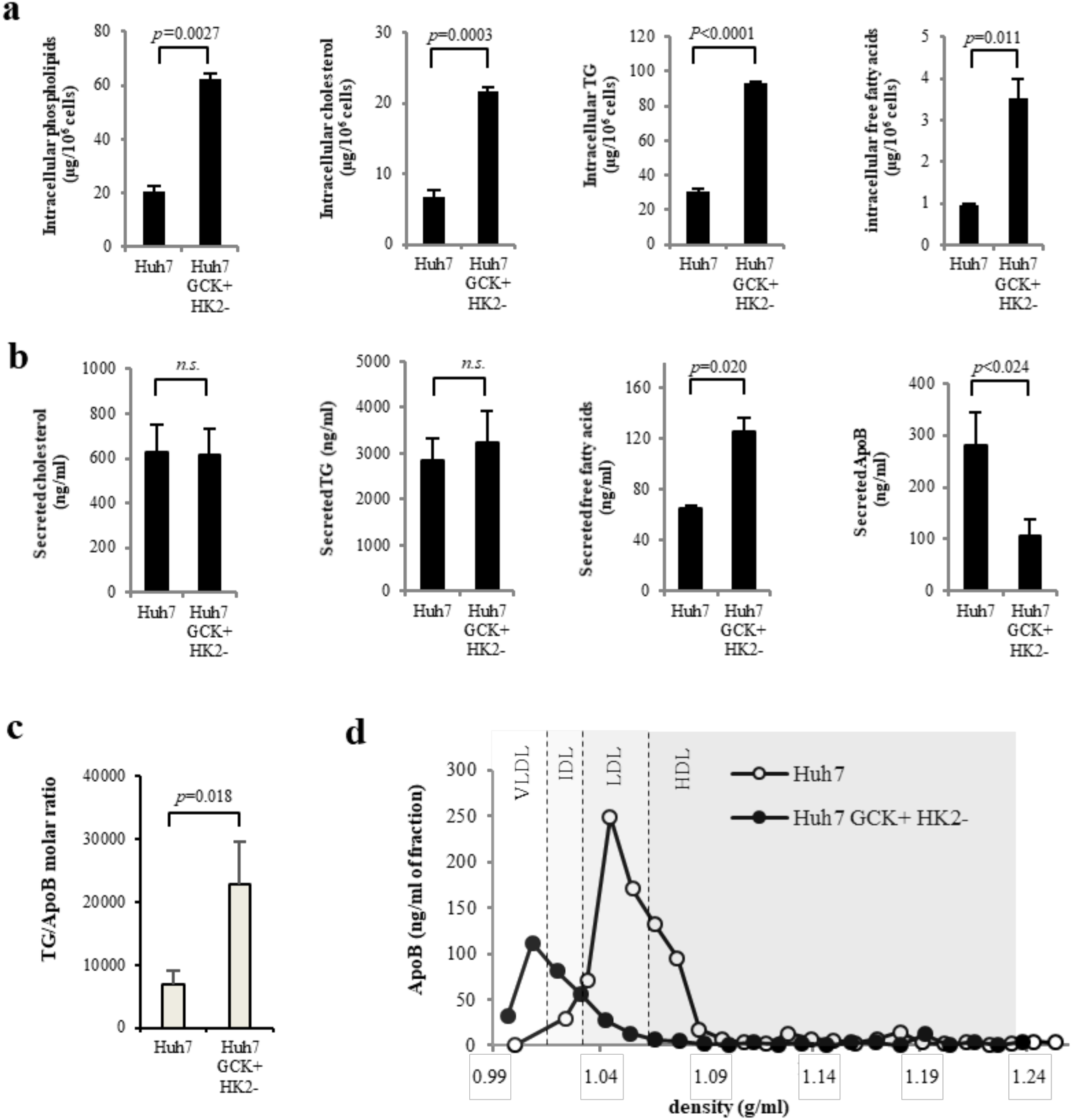
Lipogenesis and very-low-density lipoproteins (VLDL) secretion are restored in Huh7-*GCK*^*+*^/*HK2*^*−*^ cells. **a** Quantification of intracellular lipids in total cell extracts of Huh7 and Huh7-*GCK*^*+*^*HK2*^*−*^ cells. **b** Lipids and ApoB secretions in supernatants of cells cultured 24h without FCS. **c** TG/ApoB molar ratio calculated from quantifications determined in b. **d** Supernatants of Huh7 and Huh7-*GCK*^*+*^/*HK2*^*−*^ were analyzed by ultracentrifugation on iodixanol density gradients. ApoB was quantified in each fraction by ELISA. Presented data correspond to means ± SEM (n≥3, *p* value determined by Student’s t-test) except for d) that shows one representative experiment.

**Figure 5.**
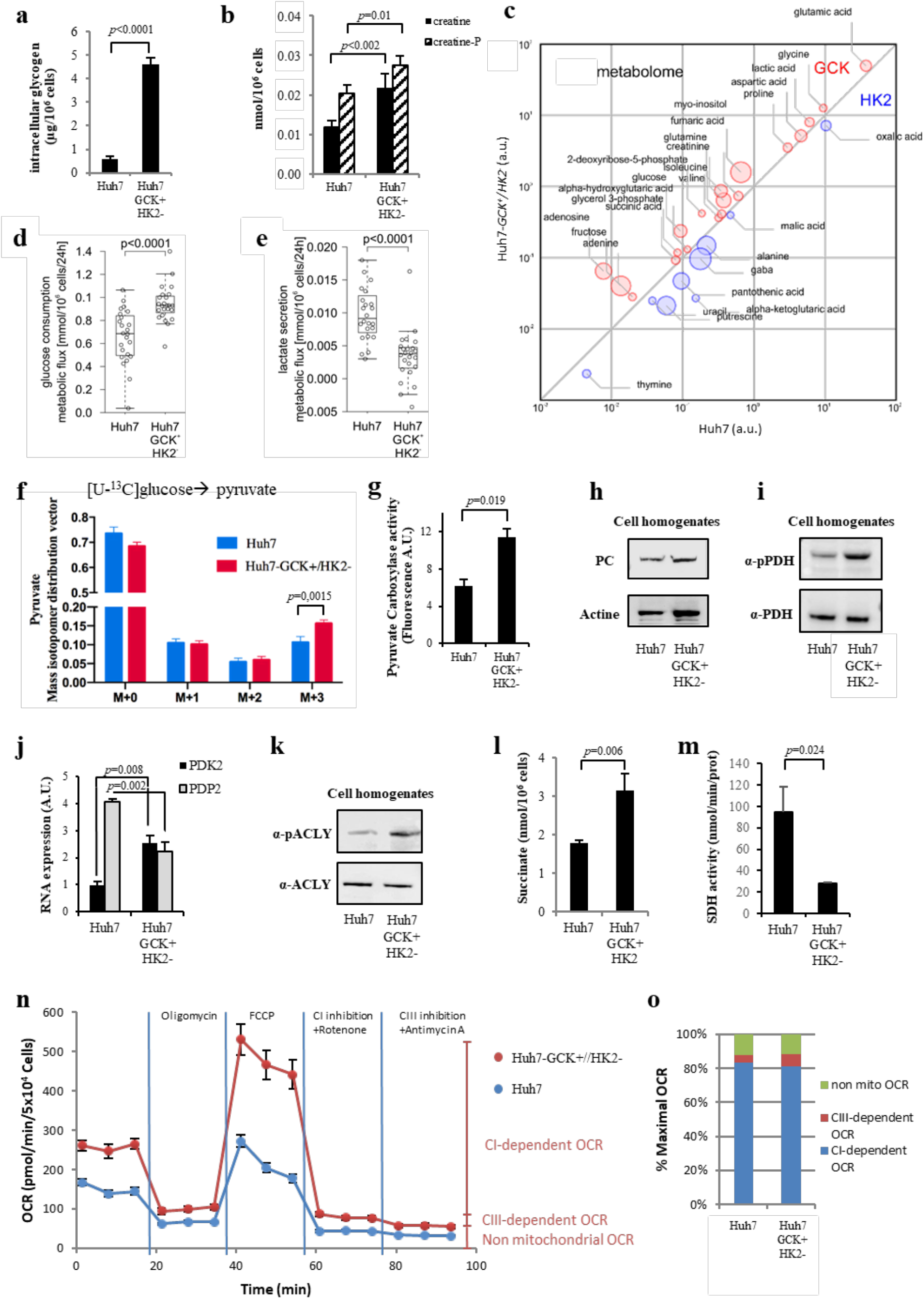
TCA rewiring after hexokinase isoenzyme switch in Huh7 cells. **a** Glycogen quantification. **b** Creatinine and creatinine-P quantification. **c** This bubble chart compares intracellular metabolomes of Huh7 and Huh7-*GCK*^*+*^/*HK2*^*−*^ cells. Metabolite pool sizes larger in Huh7 are indicated in blue, whereas the one larger in Huh7-*GCK*^*+*^/*HK2*^*−*^ are shown in red. The size of bubbles inversely scales with *p* values between 5.10^-2^ and 1.10^-17^ of differential metabolomics responses. **d, e** Metabolic fluxes for overall glucose consumption (d) and lactate secretion (e) by Huh7 and Huh7-*GCK*^*+*^/*HK2*^*−*^ cells. Indicated values correspond to differences in glucose or lactate concentrations in extracellular culture medium before and after 24h of culture. **f** Mass isotopomer distribution vector of pyruvate in cells cultured with [U-^13^C]-glucose. **g** Pyruvate carboxylase (PC) activity determined in cell homogenates. **h** Western-blot analysis of PC expression in Huh7 and Huh7-*GCK*^*+*^/*HK2*^*−*^ cells. **i**) Western-blot analysis of pyruvate dehydrogenase (PDH) E1-alpha subunit phosphorylation at Ser293. **j** RNAseq quantification of pyruvate dehydrogenase kinase 2 (PDK2) and pyruvate dehydrogenase phosphatase 2 (PDP2)(BH adjusted *p* value<0.05 from transcriptomic data). **k** Western-blot analysis of ATP-citrate Lyase (ACLY) phosphorylation at Ser455. **l** Succinate quantification in cell homogenates. **m** Succinate dehydrogenase (SDH) activity determined in cell homogenates. **n** Oxygen consumption rate (OCR) in Huh7 and Huh7-*GCK*^*+*^/*HK2*^*−*^ cells was determined with a Seahorse analyzer before and after the addition of oligomycin (Complex V inhibitor), FCCP (uncoupling agent), rotenone (Complex I inhibitor) and antimycin A (Complex III inhibitor). **o** Non-mitochondrial, complex I-dependent and complex III-dependent maximal OCR were calculated from n. Data correspond to means ± SEM (n≥3).

### Differential activity of the tricarboxylic acid cycle (TCA) in Huh7 and Huh7-*GCK*^+^/*HK2*^−^

We observed that GCK expression increased the intracellular content in lipids, resulting in accumulation of lipid droplets and secretion of VLDL. A rewiring of cellular metabolism towards energy storage in Huh7-*GCK*^*+*^/*HK2*^*−*^ was thus suspected and confirmed by the accumulation of glycogen, creatinine and creatinine-P (Fig. 5a and b), a feature of functional hepatocytes. To further determine the consequences of replacing HK2 by GCK, we quantified prominent intracellular metabolites via GC-MS. Figure 5c shows relative intracellular quantities of metabolites that are significantly different between Huh7 and Huh7-*GCK*^*+*^/*HK2*^*−*^. Among differentially represented metabolites, higher levels of glucose, glycerol-3-phosphate and lactic acid were detected in Huh7-*GCK*^*+*^/*HK2*^*−*^ cells. Several intermediates of the TCA cycle (succinic acid, fumaric acid, alpha-ketoglutaric acid), and metabolites directly connected to it (GABA, glutamic acid, glutamine, aspartic acid) were also differentially present between the two cell lines. This supports a significant modulation of central carbon metabolism at both the level of glycolysis and TCA cycle. This led to investigate glucose catabolism in further details. Glucose consumption and stable isotope incorporation from [U-^13^C]-glucose into pyruvate were both increased in Huh7-*GCK*^*+*^/*HK2*^*−*^cells compared to Huh7 cells (Fig. 5d and f). This increased glycolytic flux together with a reduced lactate secretion (Fig. 5e) is likely to account for the elevation of lactate levels and suggest that the increased pyruvate production essentially fuels mitochondrial TCA cycle in Huh7-*GCK*^*+*^/*HK2*^*−*^cells.

Pyruvate entering the mitochondria downstream of glycolysis can be either oxidized by pyruvate dehydrogenase (PDH), producing acetyl-CoA, or converted into oxaloacetate (OAA) by pyruvate carboxylase (PC). Acetyl-CoA and OAA are then combined in the TCA cycle to form citrate. *De novo* lipogenesis requires citrate egress from the TCA cycle to serve as a precursor of cytosolic acetyl-CoA for further synthesis of fatty acids. In Huh7-*GCK*^*+*^/*HK2*^*−*^ cells, we observed both an increased activity of PC (Fig. 5g) without changes in protein expression (Fig. 5h) and an increased phosphorylation of pyruvate dehydrogenase (PDH), which is indicative of a reduced activity of this enzyme (Fig. 5i). This is consistent with the increased expression of the PDH kinase PDK2 and the decreased expression of the PDH phosphatase PDP2 in Huh7-*GCK*^*+*^/*HK2*^*−*^ cells that regulate the PDH phosphorylation state (Fig. 5j). A rebalanced usage of pyruvate in Huh7-*GCK*^*+*^/*HK2*^*−*^ cells maintains a functional TCA cycle and supports lipogenesis. In Huh7-*GCK*^*+*^/*HK2*^*−*^ cells, we also observed an increased phosphorylation of ATP citrate lyase (ACLY), the first enzyme of the fatty acid synthesis pathway, indicating an enhanced activity of this enzyme (Fig. 5k). This reaction also regenerates OAA in the cytosolic compartment. Interestingly, transcriptomic data show that PCK1 which converts OAA to phosphoenolpyruvate (PEP), is overexpressed in Huh7-*GCK*^*+*^/*HK2*^*−*^cells compared to Huh7 (FC > 32).

A shift from pyruvate oxidation to carboxylation is observed in cancer cells where succinate dehydrogenase (SDH) is inactivated by mutation and OAA can only be generated through PC activity (29). SDH inhibition leads to succinate accumulation, especially in activated immune cells (30). Interestingly, higher levels of succinate and a reduced activity of SDH were measured in Huh7-*GCK*^*+*^/*HK2*^*−*^ cells compared to Huh7 (Fig. 5l and m). Even though SDH is also part of the complex II of the mitochondrial respiratory chain, we observed that the overall oxygen consumption was increased in Huh7-*GCK*^*+*^/*HK2*^*−*^ (Fig. 5n) with increased basal and maximal respiration, ATP production and spare respiration capacity (Supplementary Fig. 6). Functional analysis of the respiratory chain showed that oxygen consumption in Huh7 and Huh7-*GCK*^*+*^/*HK2*^*−*^ cells was mainly dependent on complex I which is fueling complex III (Fig. 5n and o). Thereby, the HK isoenzyme switch rewired the TCA cycle promoting carboxylation of pyruvate into OAA in the presence of a reduced SDH activity and increased respiration through complex I.

### Restored innate immune sensitivity in Huh7-*GCK*^+^/*HK2*^−^

Lipid accumulation in hepatocytes is incriminated in hepatic inflammation (31) and TCA cycle rewiring is associated with innate immunity activation (32). These two events were observed when replacing HK2 by GCK in Huh7 cells, questioning the immune status of these cells and their sensitivity to antitumor immunity. The functional analysis of gene ontology (GO) terms associated to differentially expressed transcripts revealed an enrichment in terms related to the regulation of innate immunity. The gene signature associated with type-I interferon (IFN) signaling pathway scored among the top enriched terms of upregulated genes in Huh7-*GCK*^*+*^/*HK2*^*−*^ cells (Supplementary Fig. 7). Within the 91 gene members of this GO term, 20 transcripts of Type I-IFN signaling were significantly up-regulated in Huh7-*GCK*^*+*^/*HK2*^*−*^ compared to Huh7 (Fig. 6a). This includes interferon regulatory factors (IRF1, IRF7 and IRF9), IFN-stimulated genes (ISGs) such as ISG15, MX1, OAS1, OAS3, RNaseL and signaling intermediates such as IKBKE coding for IKK∊ (Fig. 6b). The chaperon HSP90AB1, which is involved in the phosphorylation and activation of STAT1, was also induced (33). In contrast, two genes were significantly down-regulated in Huh7-*GCK*^*+*^/*HK2*^*−*^ compared to Huh7, the RNaseL inhibitor ABCE1, and TRIM6, an E3 ubiquitin-protein ligase regulating IKK∊. Among the members of the RIG-I-like receptor (RLR) family involved in immune sensing and antitumor defense (34), the expression of IFIH1 (also known as MDA5) was slightly increased in Huh7-*GCK*^*+*^/*HK2*^*−*^, while RIG-l itself remained unchanged. Such identified gene sets suggest that Huh7-*GCK*^*+*^/*HK2*^*−*^ cells may be better equipped than Huh7 cells to respond to innate immune signals. Thus, we compared the innate immune response of the two cell lines when stimulated by RIG-I or MDA5 ligands, known to sense different double-stranded RNA (dsRNA) in viral immune recognition. In order to quantify cellular activation by dsRNA, immuno-stimulatory challenge assays were conducted with triphosphate-hairpin RNA (3p-hpRNA) or polyinosinic-polycytidylic acid (poly(I:C)). Interferon-sensitive response element (ISRE)-dependent transcription was efficiently induced by these RLR ligands in Huh7-*GCK*^*+*^/*HK2*^*−*^ cells whereas Huh7 response was very limited even at high doses of immuno-stimulatory ligands (Fig. 6c).

**Figure 6.**
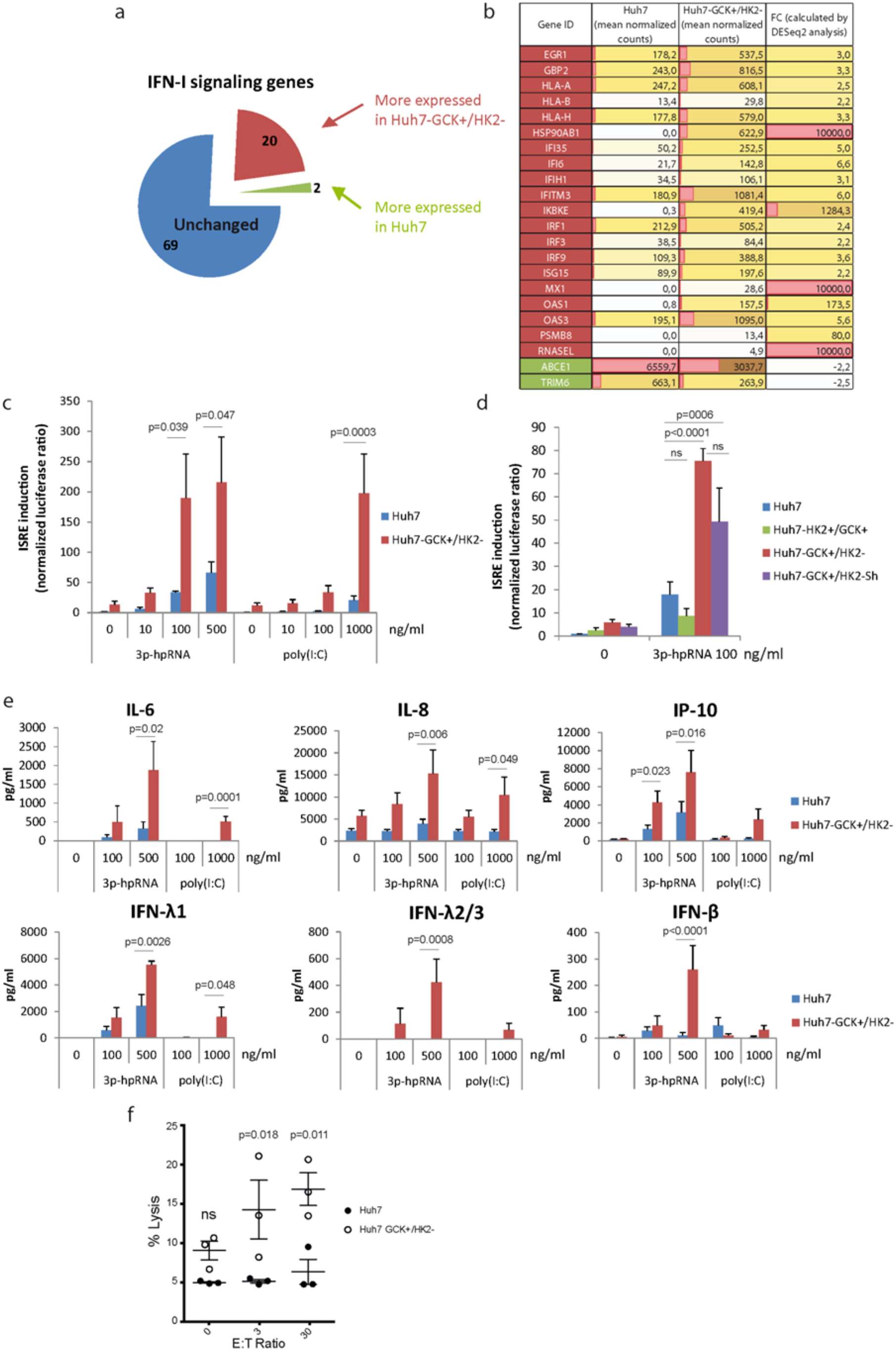
Innate immune response is enhanced in Huh7-GCK^+^/HK2^−^ cells. **a** Sector chart from the transcriptomic study showing genes included in the GO-term “Type I-IFN signaling pathway”. **b** List of genes significantly up-regulated in purple or down-regulated in green (|FC|>2, BH adjusted *p* value<0.05) in Huh7-*GCK*^*+*^/*HK2*^*−*^compared to Huh7 cells. **c-e** Cells were stimulated or not for 48h with 3p-hpRNA (RIG-I ligand) or poly(I:C) (IFIH1/MDA5 ligand). ISRE-luciferase expression was monitored and normalized to Renilla luciferase (c, d). Cell supernatants were assayed for cytokine concentration by multiplex assays (e). **f** NK cell mediated lysis of Huh7 or Huh7-*GCK*^*+*^/*HK2*^*−*^. Hepatoma cells were seeded 24h before NK cells addition for 4h at effector to target (E:T) ratio of 0, 3 or 30. After harvesting, cell lysis was determined by the percentage of PI+ cells on gated hepatocytes. *p* values were obtained from 2-way ANOVA analyses comparing matched cell means with Sidak’s correction for multiple comparison, with α=0.05.

To investigate whether this differential sensitivity to RLR ligands is linked to GCK expression or HK2 knockout, we used Huh7 cells transduced for GCK expression so that they express both HK2 and GCK (Huh7-*GCK*^*+*^/*HK2*^*+*^). In these cells, the response to RIG-I ligation did not differ from that of Huh7 cells suggesting that GCK expression alone is not sufficient to restore immune sensitivity (Fig. 6d). When HK2 expression was repressed in these cells with a shRNA (Huh7-*GCK*^*+*^/*HK2*^*−*^Sh) (see Supplementary Fig. 8 showing 95% extinction of HK2 protein), ISRE response to RIG-I signaling was restored to a level similar to that observed in Huh7-*GCK*^*+*^/*HK2*^*−*^ cells (Fig. 6d). This is pointing towards HK2 as a negative regulator of RLR signaling in HCC cells and suggests that the *GCK*-to-*HK2* isoenzyme switch during malignant transformation of hepatocytes is accompanied by a reduced sensitivity to innate immune signals. The higher sensitivity to RLR ligands of Huh7-*GCK*^*+*^/*HK2*^*−*^ cells also resulted in increased secretion of inflammatory interleukins (IL-6 and IL-8), antiviral cytokines (IFN-λ_1_, IFN-λ_2/3_, IFN-β), and IP-10 (Fig. 6e), indicating that both NF-κB- and IRF3-dependent signaling pathways were induced. IL-1β, TNFα, IL-12p70, GM-CSF, IL-10 and IFNγ were not detected in the supernatants of none of the cell lines, whether they were stimulated or not.

As natural killer (NK) cell-mediated lysis of tumor cells is crucial for the anti-cancer immune defense, we compared the susceptibility of the two cell lines to NK cells cytotoxicity. Figure 6f shows that Huh7 cells are resistant to NK cell-mediated lysis in contrast to Huh7-*GCK*^*+*^/*HK2*^*−*^. Thus, replacing HK2 by GCK restored Huh7 sensitivity to NK cell-mediated lysis. Similar results were obtained when NK cells were pre-activated with IL-2 (Supplementary Fig. 9). Altogether, these results demonstrate that HCC cells expressing HK2 instead of GCK exhibit an impaired response to immune signals and also a strong resistance to NK cells. Significantly, these two observations are in line with clinical data showing that elevated GCK expression is associated with prolonged survival, while elevated HK2 expression coinciding with GCK reduction correlates with shorter overall survival (Fig. 1).

## Discussion

Metabolic network rewiring is a hallmark of cancer although for most tumors, mechanisms at the origin of this metabolic reprogramming have not been elucidated. While GCK, but not HK2, is expressed in normal hepatocytes, the expression of HK2 occurs during cirrhosis and increases as the disease progresses to carcinoma. Several signaling pathways such as hypoxia inducible factors (HIF), peroxisome proliferator-activated receptors (PPAR) and phosphatidylinositol-4,5-bisphosphate 3-kinase (PI3K) might contribute to HK2 induction in fatty liver disease and its evolution towards cirrhosis and carcinogenesis (35–37). Consequently HK2 induction has been proposed as a risk marker of HCC development (16). Analyzing TCGA data from human HCC tumors, we observed that not only high levels of *HK2* but also low levels of *GCK* are of poor prognosis. In contrast, neither *HK1* nor *HK3* expression levels were correlated with survival of HCC patients. *GCK* expression is very low or not detected in biopsies from a majority of patients (65.8% of patients show RSEM values <10), whereas *HK2* is widely expressed (16) (only 5.8% of patients show RSEM values <10). This probably explains that HK2 expression is a better prognostic marker than GCK for HCC. However, when GCK and HK2 expression were combined into a single ratio, this prognostic marker outperformed HK2 or GCK expression alone. This suggests that both HK2 induction and GCK loss play a role in HCC progression. As HK2 and GCK expression tend to be mutually exclusive, both HK2 induction and GCK downregulation might have consequences on the metabolic reprogramming during malignant transformation of hepatocytes. To compare the functional consequences of the HK isoenzyme switch in HCC, we therefore expressed GCK in the reference HCC cell line Huh7 and knocked-down HK2 expression. Our comparative transcriptomic, metabolic and functional studies demonstrate that the replacement of HK2 by GCK not only restored some essential metabolic functions of normal hepatocytes such as lipogenesis, VLDL secretion and glycogen storage but also reactivated innate immune responses and sensitivity to NK-mediated cell lysis.

HCC cell lines predominantly secrete LDL-like particles, unlike normal hepatocytes, which secrete VLDL. Lipid loading of Huh7 cells with oleic acid can boost the secretion of ApoB^+^ particles but does not induce a shift from LDL to VLDL density, indicating that intracellular fatty acid accumulation of exogenous origin cannot rescue VLDL production (28). Here we show that replacing HK2 by GCK in Huh7 cells restored *de novo* fatty acid synthesis, allowing VLDL assembly/secretion in the absence of exogenous lipids. To our knowledge Huh7-*GCK*^*+*^/*HK2*^*−*^ is the first human cell model with a functional VLDL secretion pathway. Such a tool will strongly benefit the field of cardiovascular diseases and hepatic steatosis.

*De novo* fatty acid synthesis from carbohydrates requires an adequate supply in metabolic substrates, especially citrate that is produced by the TCA cycle from incoming pyruvate. The glycolytic entry point into the TCA cycle is controlled by PDH and PC that convert pyruvate into acetyl-CoA or OAA, respectively. Our data revealed that in addition to the increased production of pyruvate from glucose, PC activity is increased whereas PDH is inhibited. This suggests that pyruvate metabolism is rebalanced in favor of OAA in Huh7-*GCK*^*+*^/*HK2*^*−*^ cells, as described in healthy liver. Such a mechanism of anaplerosis is known to replenish TCA cycle intermediates and compensate citrate export out of the mitochondria for lipogenesis fueling. Increased PC activity is observed in both normal and pathological situations, mainly as a result of an increased transcription of the PC gene. In our model, mRNA and protein levels were not affected, indicating that PC activity can be regulated by alternative mechanisms depending on HK isoenzyme expression. This may relate to lower levels of oxalate, a known inhibitor of PC activity, in Huh7-*GCK*^*+*^/*HK2*^*−*^ cells (Fig. 5c and Fig. 7 discussed below).

**Figure 7.**
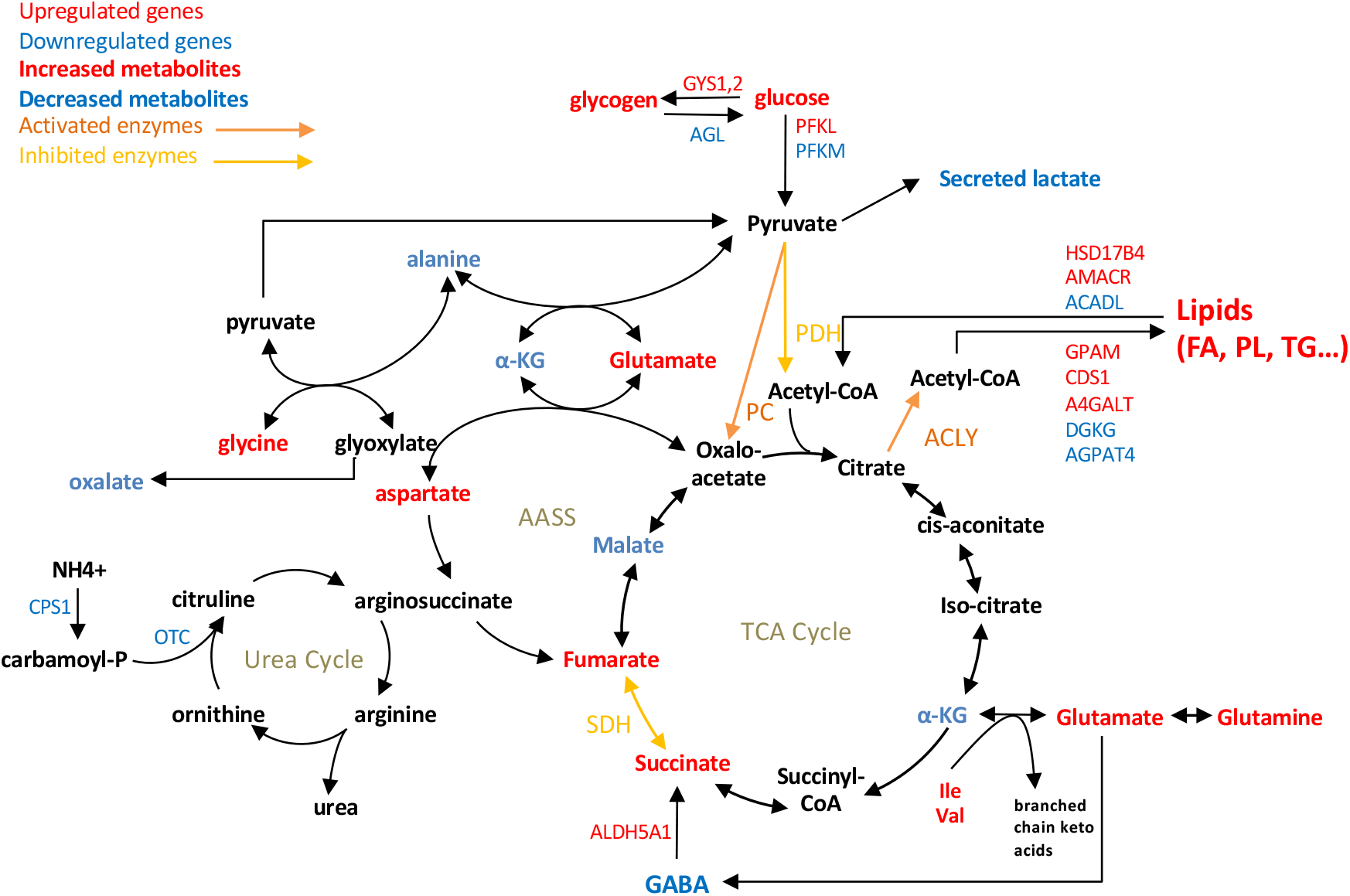
Simplified scheme of central carbon metabolism and connected pathways showing differences between Huh7-*GCK*^*+*^/*HK2*^*−*^ vs Huh7. Highlighted metabolites, enzymatic activities, and metabolism-associated genes were selected from transcriptomic (Fig. 2e), metabolomic (Fig. 5c) and enzymatic analyses (Fig. 5g-m).

A rebalanced pyruvate usage in favor of OAA is also described for instance in SDH-deficient neuroendocrine tumor cells, where succinate accumulates and PC activity is increased to maintain OAA production, replenish the oxidative TCA cycle and support aspartate synthesis (29). Interestingly, in comparison to Huh7 cells, succinate and aspartate levels are elevated in Huh7-*GCK*^*+*^/*HK2*^*−*^ where SDH activity is reduced, suggesting a direct link between PC and SDH activity in hepatocytes. Several mechanisms inhibiting SDH have been described (38). Modification of the expression of SDH subunits is unlikely as no variation was observed at the transcriptomic level. Itaconate is a weak inhibitor of SDH produced by IRG1 from aconitate but this metabolite was not detected and IRG1 mRNA was absent from the transcriptome of both cell lines. Whether fumarate or other metabolites are responsible for the reduced SDH activity in GCK-expressing cells remains to be investigated. Finally, SDH-deficient cells and LPS-stimulated macrophages have been shown to elicit a hypoxic-like phenotype through accumulation of large amounts of succinate and stabilization of HIF-1α (39, 40). Despite an elevated succinate steady-state level in Huh7-*GCK*^*+*^/*HK2*^*−*^ compared to Huh7 cells, we observed no difference in HIF-1α stabilization neither at basal level nor upon induction (Supplementary Fig. 10). This suggested that the reduction of SDH activity in Huh7-*GCK*^*+*^/*HK2*^*−*^ cells was not strong enough to induce such a pseudo-hypoxic phenotype.

Our gene-centric metabolic analysis of transcriptomic data revealed a wide spreading of metabolic modifications resulting from HK isoenzyme switch. Illustrating these modifications, Figure 7 is an attempt to integrate the observed changes in central carbon metabolism and closely connected metabolic pathways. In particular, decreased level of alanine and increased aspartate concentration in Huh7-*GCK*^*+*^/*HK2*^*−*^ cells could be an indirect effect of PC activation that uses pyruvate for the synthesis of OAA. As a consequence, hepatic transaminases may balance intracellular pools of OAA, aspartate, alanine and pyruvate. Glutamate and GABA levels were also modified, thus supporting anaplerosis of the TCA cycle through glutamine consumption and the GABA shunt pathway, respectively. We also observed lower levels of oxalate, an end-product of glyoxylate degradation. In Huh7-*GCK*^*+*^/*HK2*^*−*^ cells, increased levels of the enzyme alanine-glyoxylate and serine-pyruvate aminotransferase (AGXT) could account for this phenotype as it converts alanine and glyoxylate into pyruvate and glycine, which is also increased. Interestingly, high level of AGXT is a good prognostic marker for HCC (41). Consistently, it was found that oxalate inhibits liver PC, resulting in reduced gluconeogenesis and lipogenesis (42, 43). Thus, a higher PC activity could be explained by lower levels of oxalate in Huh7-*GCK*^*+*^/*HK2*^*−*^ cells. We also observed that isoleucine and valine levels increased while BCAT1 (branched chain amino acid transaminase 1) predominant transcripts decreased. This suggests a reduced catabolism of branched chain amino acids in Huh7-*GCK*^*+*^/*HK2*^*−*^ cells. Again, low levels of BCAT1 is a good prognostic marker for HCC and oral supplementation with branched chain amino acids has been shown to reduce the risk of liver cancer in cirrhotic patients (44, 45). If some metabolic modifications seem to advocate for the restoration of a normal hepatocyte phenotype following the replacement of HK2 by GCK, it cannot be a general statement. Indeed, the urea cycle was also impacted in Huh7-*GCK*^*+*^/*HK2*^*−*^ cells with lower levels of CPS1 and OTC, which are also observed in aggressive HCC tumors (46). Altogether, our results demonstrate the broad impact of replacing HK2 by GCK in HCC cells, and the key role played by the HK isoenzyme switch in HCC tumor metabolism.

We discovered that HK isoenzyme expression not only controls hepatic metabolic functions but also interferes with intrinsic innate immunity of hepatocytes and antitumor immune surveillance. Several reports have recently established functional links between glucose metabolism and signaling pathways downstream of innate immunity sensors of the RLR family, RIG-I and MDA5 (47–49), which are usually associated with antiviral responses. However, RIG-I expression is downregulated in hepatic cancer tissues and low RIG-I expression is correlated with poor survival of patients, whereas RIG-I expression in HCC cell lines enhanced IFN response and cancer cell apoptosis (34). This suggests an unexpected role of this receptor family in the antitumor response. Here we show that Huh7 cells expressing GCK instead of HK2 exhibit a higher sensitivity to RIG-I and MDA5 ligands, and produce higher levels of type I/III IFNs and inflammatory cytokines. This immune phenotype occurs in a context of reduced SDH activity and increased intracellular content in succinate (Fig. 5l-m). A pro-inflammatory function of immune cells such as macrophages was previously linked to TCA rewiring, with reduced SDH activity resulting in succinate accumulation (39, 50, 51). Succinate can also be secreted from LPS-activated macrophages and activate its cognate receptor, succinate receptor 1 (SUCNR1, previously known as GPR91) in an autocrine and paracrine manner to further enhance production of IL-1β (52). Interestingly, glucose metabolism promotes RIG-I and MDA5 signaling through the O-GlcNAcylation of the mitochondrial adaptor MAVS (47). Thus, an intriguing hypothesis is that GCK expression could facilitate MAVS signaling by increasing UDP-GlcNAc through upregulation of the hexosamine biosynthetic pathway. HK2 binding at the surface of mitochondria may also compete with pyruvate carboxylase, metabolites or mitochondria factors known to control MAVS signaling (47–49). Here we show that HK2 knockdown promotes RIG-I-induced ISRE-dependent transcription (Fig. 6d). This is consistent with the results obtained by Zhang W. et al. (48), indicating that HK2 interaction with MAVS restrains RIG-I-induced IFN-β secretion. Further investigations are now required to decipher the molecular links between metabolism and immune responses.

Beyond the inhibition of RLR signaling, other mechanisms might contribute to tumor escape from immune surveillance in HCC patients. In advanced-stage HCC patients, NK cells often exhibit reduced infiltration and impaired functional activities (53). We thus compared the sensitivity of Huh7-*GCK*^*+*^/*HK2*^*−*^ cells to Huh7, and found that sensitivity to NK cell lysis is restored when HK2 is replaced by GCK. When analyzing cell surface expression of the NK cells inhibitors, HLA class I and MICA/B, no significant changes were observed between cell lines (Supplementary Fig. 9). In contrast, an increased transcription and surface expression of ICAM1 (FC=2.6; Supplementary Fig. 9) was observed in Huh7-*GCK*^*+*^/*HK2*^*−*^. Since ICAM-1 binding to active LFA-1 at the surface of NK cells is essential for granule polarization and efficient killing of the target cells (54), its enhanced exposition at the surface of Huh7-*GCK*^*+*^/*HK2*^*−*^ may contribute to their higher sensitivity to NK cell-mediated killing. These results suggest that HK2 expression at the expense of GCK in HCC tumors decreases immune responsiveness and sensitivity to NK cytotoxicity, thus favoring immune escape.

Taken together, our data demonstrate that beyond glycolysis, the hexokinase isoenzyme switch in an HCC model rewires central carbon metabolism, promotes lipogenesis, enhances innate immune functions, and restores sensitivity to natural killer cells.

## Materials and Methods

### Materials

Unless otherwise specified, chemicals were from Merck Sigma-Aldrich. The RIG-I specific ligand 3p-hpRNA and the MDA5/TLR3 ligand poly(I:C) HMW (High Molecular Weight) were from Invivogen.

### Cell cultures

Huh7 cells and derivatives were grown as previously described (55) in DMEM, 10% fetal calf serum (FCS), penicillin/streptomycin, 1 mM pyruvate, 2 mM L-glutamine. Culture medium and additives were from Gibco except FCS (Dominique Dutcher).

### Cell lines

15×10^4^ Huh7 cells were transduced for GCK expression at different multiplicities of infection (lentiviral transduction using the pLEX-GCK construct). The Huh7-*GCK*^*+*^/*HK2*^*+*^ cells were then cultured for 7 days with puromycin (1 μg/mL) before amplification. HK2 knock-out was achieved using the CRISPR/Cas9 system as previously described (56) to obtain Huh7-*GCK*^*+*^/*HK2*^*−*^ cells. A single guide RNA (sgRNA) pair was designed for double nicking using the CRISPR Design Tool (http://tools.genome-engineering.org). The guide sequence oligos (sgRNA_1_(*HK2*): 5’-CACCGTGACCACATTGCCGAATGCC-3’ and sgRNA_2_(*HK2*): 5’-CACCGTTACCTCGTCTAGTTTAGTC-3’) were cloned into a plasmid containing sequences for Cas9 expression and the sgRNA scaffold (pSpCas9(BB)-2A-GFP, Addgene plasmid #48138). 48 h post-transfection, cells were sorted by FACS based on the transient expression of GFP and cloned by limiting dilution. Effective deletion of HK2 was assessed by qPCR.

For *HK2* knock-down, Huh7-*GCK*^*+*^/*>HK2*^*+*^ cells were transduced with lentiviral vectors expressing HK2-targeting shRNAs, and antibiotic selection was applied (hygromycin; 100 μg/ml). The *HK2*-targeting sequence 5’-CCGGCCAGAAGACATTAGAGCATCTCTCGAGAGATGCTCTAATGTCTTCTGGTTTTTT-3’ was cloned in the pLKO.1 hygro vector (a gift from Bob Weinberg; Addgene plasmid #24150). HK2 expression in Huh7-*GCK*^*+*^/*HK2*^*+*^ and Huh7-*GCK*^*+*^/*HK2*^*−*^*Sh* was analyzed on cell lysates by western blotting (Supplementary Fig. 8).

### Enzymatic activity assays

Cells were trypsinized, washed twice, and cell pellets were stored at −80°C. Protein extractions and assays were performed as previously described for hexokinase (57–59) and pyruvate carboxylase (60). Specific enzymatic assays are described in Supplementary Methods.

### Metabolomics profiling

Cells were seeded at 13×10^5^ cells in 75cm^2^ dishes. After 24 h, culture medium was replaced and cells were further incubated during 24 h. Culture supernatants were collected and stored at −80°C. Cells were harvested, washed twice with ice-cold PBS, and cell pellets were stored at −80°C. To analyze catabolic glucose flux, cells were incubated with [U-^13^C]-glucose (Sigma-Aldrich; 389374-2G) at 12.5 mM and unlabeled glucose at 12.5 mM for 24h. Extraction, separation and analysis of the metabolites by GC-MS are described in Supplementary Methods.

### Transcriptome profiling of Huh7 and Huh7-*GCK*^*+*^/*HK2*^*−*^ cell lines

Transcriptome profiling was performed by next generation sequencing (Profilexpert, Lyon). To identify differentially expressed genes, raw data were processed using the DESeq2 analysis pipeline (61). See Supporting Information and Gene Expression Omnibus database with the accession number GSE144214 for entire raw data.

### Pathway analysis

The list of transcripts differentially expressed in Huh7 and Huh7-*GCK*^*+*^/*HK2*^*−*^ cell lines was analyzed by gene set enrichment analysis (IPA, Build version: 486617M, Qiagen) weighted by their corresponding fold-change and *p* value. The fold-change cut-off of mean expression for each transcript was set at 2 with an adjusted *p* value<0.05. The list of genes associated with “Type I-IFN signaling pathway” was defined in the AmiGO 2 database. Expression data of these genes were retrieved from the transcriptomes of Huh7-*GCK*^*+*^/*HK2*^*−*^ and Huh7, and correspond for each gene to the most differentially expressed transcript.

### Cell migration assay

2×10^4^ cells were plated in the upper chamber of transwells (Sarstedt, PET 8.0-μm, TL - 833932800) with DMEM without FCS to allow migration for 24 h at 37°C. DMEM with 10% FCS was distributed in each well, below the chamber. Chambers were gently picked up before a brief PBS rinse and 0.05% crystal violet coloration. The migrated cells were analyzed a Leica M50 microscope using a magnification factor of 20x. The number of cells that have migrated through the membrane and attached on the underside of the membrane were counted using the software Image J.

### Respiration Assay

Respiration was measured using an Extracellular Flux Analyzer (Seahorse Bioscience). The assay was performed according to the Agilent Seahorse XF Cell Mito Stress Test as detailed in Supplementary Methods. The number of cells was determined at the end of the run using the SRB assay as previously described (62) or in plate cell counting by Cytation 1 imaging reader (Biotek).

### Intracellular lipid staining

For microscopic observations, cells were fixed 48 h post-seeding, and intracellular lipids stained with Oil-Red-O as described in Supplementary Methods. For the quantification of intracellular lipid droplets by FACS, cells were stained with the 4,4-difluoro-1,3,5,7,8-Pentamethy-4-Bora-3a,4a-Diaza-s-indacene (BODIPY) 493/503 dye (Tocris Bio-Techne).

### Protein, ApoB, and Lipid Quantification

Protein concentration was determined using the DC Protein Assay (Bio-Rad). ApoB concentration in medium and gradients fractions was determined by ELISA as previously described (63). Total concentrations of cholesterol, phospholipids, and triglycerides (TG) were determined using specific assays from Millipore Sigma-Aldrich (ref. MAK043, MAK122 and MAK266 respectively). Free Fatty Acids were quantified using a specific assay kit from Abcam (ref. ab65341).

### Iodixanol density gradients

Iodixanol gradients were prepared as previously described (64). One ml of culture supernatant was applied to the top of 6 to 56% iodixanol gradients and centrifuged for 10h at 41,000rpm and 4°C in a SW41 rotor. The gradient was harvested by tube puncture from the bottom and collected into 22 fractions (0.5 ml each). The density of each fraction was determined by weighing.

### Metabolic network coherence computational analysis

In order to measure the consistency of differentially expressed genes with a metabolic network, we employed the metabolic network coherence measure introduced by Sonnenschein *et al.* (65) (see Supplementary Methods for details). This approach was previously applied to various disease-related transcriptome profiles (66, 67) and for extracting information on the genetic control of metabolic organization (68). Recently, detailed theoretical analysis of the extended version of the method has been performed by Nyczka and Hütt (69)

### Western-Blot Analysis

Protein expression was determined by standard western-blot analysis (See Supplementary Methods for details).

### RLR stimulation

Cells were seeded in 96-well or 24-well plates. After 24 h, cells were co-transfected with indicated doses of the RIG-I ligand 3p-hpRNA or the MDA5/TLR3 ligand poly(I:C) HMW together with the pISRE-luc (1.25 μg/ml) and pRL-SV40 (0.125 μg/ml) reporter plasmids using the JetPEI-Hepatocyte reagent (Polyplus Transfection). Manufacturer’s instructions were followed. After 48 h, supernatants were collected for cytokine quantification. Firefly and Renilla luciferase expressions within cells were determined using the Dual-Glo luciferase Assay system (Promega) and an Infinite M200 microplate reader (TECAN).

### Cytokine assays

Clarified culture supernatants were collected and stored at −20°C. IL-8 was quantified using the Cytometric Bead Array for human IL-8 (BD Biosciences). Other cytokines were assayed using the LEGENDPlex multiplex assay (Human Anti-Virus Response Panel, BioLegend). Fluorescence was analyzed using a FACS Canto II (BD Biosciences).

### NK cell cytotoxicity test

NK cells were isolated as described in Supplementary Methods from human buffy coats of healthy donors obtained from the Etablissement Français du Sang. Informed consent was obtained from donors and experimental procedures were approved by the local institutional review committee. Huh7 or Huh7-*GCK*^*+*^/*HK2*^*−*^ were seeded at 1×10^5^ cells per well in a 24-well plate in RPMI-1640 (Gibco) with 10% FCS and 40 μg/ml gentamycin. After 24 h, 3×10^5^ or 3×10^6^ NK cells were added to the culture wells. The cytotoxicity assay was performed for 4 h at 37°C, under 5% CO_2_. Target hepatoma cells were harvested after trypsination, labelled with propidium iodide (PI) and analyzed by FACS. Cell death was monitored after morphological gating on hepatocytes.

### Statistics and reproducibility

All the statistical analyses were performed with GraphPad Prism or Analyse-it softwares. Details of statistical analyses can be found in figure legends.

## Acknowledgments

We acknowledge the contribution of the Genomics and Microgenomics platform ProfileXpert (University Lyon 1, SFR santé LYON-EST, UCBL-Inserm US 7-CNRS UMS3453) and SFR Biosciences (UMS3444/CNRS, US8/Inserm, ENS de Lyon, UCBL) facilities: AniRA-Cytometry, AniRA-ImmOs metabolic phenotyping and LYMIC-PLATIM microscopy. We gratefully thank Laurence Canaple for technical assistance. This work was supported by the Fondation pour la Recherche Médicale (FRM), grant number DEQ20160334893 to VL. F.V.F. is grateful for the support by grants CA154887, GM115293, CRN-17-427258, NSF GRFP, and the Science Alliance on Precision Medicine and Cancer Prevention by the German Federal Foreign Office, implemented by the Goethe-Institute, Washington, DC, USA, and supported by the Federation of German Industries (BDI), Berlin, Germany.

## Author Contributions

L.P-C., P-O.V., O.D., V.L. designed the experiments with critical advices from G.J.P.R., P.A., R.R. and F.V.F.; C.J., A.A-G., K.O., B.P., G.J.P.R., N.A, R.R. and F.V.F. performed experiments and analyzed the data; P.N. and M-T.H. performed metabolic network computational analysis; L.P-C., P-O.V., P.A., F.V.F, V.L. and O.D. analyzed the data, prepared figures and wrote the manuscript.

