## Supplementary Figures for "Lipogenesis and innate immunity in hepatocellular carcinoma cells reprogrammed by an isoenzyme switch of hexokinases"

**This PDF file includes:**

Supplementary Methods
Supplementary Figures 1 to 10

**Other supplementary materials for this manuscript include the following:**

Supplementary Tables 1 to 3

### Supplementary Methods

#### Hexokinase activity assay.

The method used for monitoring HK activity in cells lysates was adapted from *Kuang et al.*, as previously described (1–3). Cellular pellets stored at -80°C were thawed and immediately homogenized ($2x10^6$ cells/100 µl) in precooled reaction buffer. (0.05 M Tris–HCl, 0.25 M sucrose, 0.005 M EDTA, 0.005 M 2-mercaptoethanol, pH=7.4). After 20 min incubation on ice, homogenates were pulse-sonicated 15 s at half power (EpiShear Probe Sonicator). Homogenates were then centrifuged at 500 g for 20 min at 4°C. Supernatants were immediately used for determination of HK activity, which was measured spectrophotometrically through NADP+ reduction in the glucose 6-phosphate dehydrogenase–coupled reaction. HK activity was assayed in medium containing 50 mM triethanolamine (pH=7.6), 10 mM $MgCl_2$, 1.4 mM $NADP^+$, with variable concentration of glucose and 1 U glucose 6-phosphate dehydrogenase (S. cerevisiae), equilibrated to 37°C. The reaction was started by addition of ATP (final concentration 1.9 mM), and absorbance was continuously recorded for 30 min at 340 nm (TECAN Infinite M200).

#### Pyruvate Carboxylase activity assay.

The method used for quantification of PC activity was adapted from Payne et al. (4). Briefly, cells were centrifuged, washed twice with ice-cold PBS before homogenization in Tris-HCL 100 mM, pH=8.0 using a Dounce homogenizer. Homogenates were pulse-sonicated 15 s at half power (EpiShear Probe Sonicator) before centrifugation at 500 g for 5 min. Supernatants were immediately used for the assay. PC activity was assayed in medium containing 100 mM Tris-HCl, 50 mM $NaHCO_3$, 5 mM $MgCl_2$, 0.1 mM Acetyl-CoA, 0.25 mM 6,6′-Dinitro-3,3′-dithiodibenzoic acid (DTNB), 5 mM ATP, 5 mM pyruvate, citrate synthase and cofactors. Reduction of DTNB by the generated free CoA was measured continuously by Abs at 412 nm and recorded for 30 min (TECAN Infinite M200). The same assay was performed in absence of pyruvate to subtract background signal.

#### Metabolomics profiling of HCC model cells.

For metabolomics profiling, cells were seeded at $13x10^5$ cells per 75 $cm^2$ dishes. After 24 h, supernatant was removed and replaced by fresh culture medium. For quantification of metabolic flux from glucose, culture medium was supplemented with both [U-$^{13}$C]-glucose (Sigma-Aldrich; 389374-2G) and unlabeled glucose at a 50:50 ratio. After 24 h, cells were harvested, washed twice with ice-cold PBS and cell pellets were frozen at -80°C until metabolites extraction. Cell pellets were transferred into a pre-chilled microcentrifuge tube with 1 mL cold extraction buffer consisting of 50% methanol (A452, Fisher Scientific) in ultrapure water. Samples were then frozen in liquid nitrogen, thawed, and placed in a shaking dry bath (Thermo Fisher Scientific, Waltham, MA) set to 1100 rpm for 15 min at 4°C. After centrifugation for 15 min at 12500 g and 4°C (Sorvall, Thermo Fisher Scientific) using a fixed-angle F21-48x1.5 rotor, supernatants were collected and dried by vacuum centrifugation overnight. Dried metabolites were derivatized by addition of 20 µL of 2.0% methoxyamine-hydrochloride in pyridine (MOX, TS-45950, Thermo Fisher Scientific) followed by incubation during 90 min in shaking dry bath at 30°C and 1100 rpm. Ninety µL of N-methyl-N-trimethylsilyltrifluoroacetamide (MSTFA, 701270.201, Macherey-Nagel) was added, and samples were incubated and shaken at 37°C for 30 min before centrifugation for 5 min at 14,000 rpm and 4°C. Metabolites contained in the supernatant were then separated by gas chromatography (GC, TRACE 1310, Thermo Fisher Scientific) coupled to a triple-quadrupole mass spectrometry system for analysis (QQQ GCMS, TSQ8000EI, TSQ8140403, Thermo Fisher Scientific), equipped with a 0.25 mm inner diameter, 0.25 µm film thickness, 30 m length 5% diphenyl / 95% dimethyl polysiloxane capillary column (OPTIMA 5 MS Accent, 725820.30, Macherey-Nagel) and run under electron ionization at 70 eV. Using established separation methods (5–7), the GC was programed with an injection temperature of 250.0°C and splitless injection volume of 1.0 µL. The GC oven temperature program started at 50°C (323.15 K) for 1 min, rising at 10 K/min to 300.0°C (573.15 K) with a final hold at this temperature for 6 min. The GC flow rate with helium carrier gas (HE, HE 5.0UHP, Praxair) was 1.2 mL/min. The transfer line temperature was set at 290.0°C and ion source temperature at 295.0°C. A range of 50-600 m/z was scanned with a scan time of 0.25 s.

**Metabolomics data processing.**
Metabolites were identified using TraceFinder (v3.3, Thermo Fisher Scientific) based on libraries of metabolite retention times and fragmentation patterns (Metaflux, Merced, CA). Identified metabolites were quantified using the selected ion count peak area for specific mass ions, and standard curves generated from reference standards run in parallel. Peak intensities were median normalized. The mean and standard deviation for each quantified metabolite was calculated for each cell line and treatment condition. A univariate t-test was used to compare treatment conditions for each metabolite and cell line.

**Transcriptome profiling**
Total RNA was extracted and purified from cell pellets using Direct-zol RNA purification kit (Zymo Research). 700 ng of total RNA were amplified (NextFlex Rapid Directional mRNA-Seq, PerkinElmer) to generate mRNA-seq libraries. Then, gene expression was analyzed by Next-Generation Sequencing (NGS) using Illumina NextSeq500. Reads were mapped on the reference genome Homo sapiens GRCh37/hg19. Raw data were processed using the DESeq2 pipeline (8) to identify differentially expressed genes.

**Metabolic network coherence computational analysis.**
We first extracted a gene-centric metabolic network from a given genome-scale metabolic model. This was achieved via the stoichiometric matrix and the gene-reaction associations contained in the metabolic model. We constructed the two projections of the bipartite graph represented by the stoichiometric matrix, yielding a metabolite-centric and a reaction-centric graph. The metabolite-centric graph allowed us to identify high-degree nodes ('currency metabolites' like $H_2O$, ATP, etc.), which are not informative about the network-like organization of the metabolic systems and need to be eliminated before interpreting the network architecture (see ref (9) and (10) for details). The degree of a node is the number of neighbors the node has in the network. The percentage of remaining metabolites is one of the parameters of our analysis. Typical values are 90 to 98 percent (i.e. a removal of the highest 2 to 10% of metabolites with the highest degree as currency metabolites). After recomputing the reaction-centric graph based on the reduced number of metabolites (Supplementary Fig. 2), we can now evaluate the gene-reaction associations to arrive at a gene-centric metabolic network (Supplementary Fig. 2). Given a set S of differentially expressed genes and the gene-centric metabolic network G, we can now analyze the subgraph of G spanned by all genes in S. The average clustering coefficient C in these subgraphs serves as a measure of the connectivity of this subgraph. The metabolic network coherence MC is the z-score of C computed with respect to a null model of randomly drawn gene sets with the same size as S (Supplementary Fig.3). In this way, MC has an intuitive interpretation: The value of MC indicates, how many standard deviations away from randomness the clustering of the subgraph spanned by the observed gene set S actually is (Supplementary Fig.3 and ref. (11)). The genome-scale metabolic models employed here are the generic human metabolic model Recon 2 (12). In general, different network measures can be used for evaluation of MC. In the scope of this study, we have tested several of them, but opted for average clustering coefficient C, as it yielded strongest statistical signal.

**Respiration Assay.**
Twenty-four hours prior measuring respiration in the Seahorse Extracellular Flux Analyzer, the cells were seeded in XF 24-wells cell culture microplates (Seahorse Bioscience) at 5x104 cells/well in 100 µL of DMEM medium supplemented with 10% FCS, 1 mM pyruvate, 2 mM L-glutamine, penicillin/streptomycin, and then incubated at 37°C / 5% CO2 during 5 h for cell attachment. Medium volume was adjusted to 250µL and cells incubated overnight. The assay was initiated by removing the growth medium from each well and replacing it with 500 µL of Seahorse assay medium (XF DMEM pH7.4 + 10 mM Glucose, 2 mM Glutamine and 1 mM sodium pyruvate) prewarmed at 37°C. Cells were incubated at 37°C for 1h to allow media temperature and pH to reach equilibrium before the first rate measurement. The oxygen consumption rate (OCR) was measured using the following Seahorse running program: injection Port A – 1.5 µM Oligomycin; Injection Port B – 0.5 µM FCCP and injection Port C – 0.5 µM Rotenone; injection port D – 0.5 µM

Antimycin A. The number of cells was determined at the end of the run after Hoechst staining and cell counting using Cytation 1 imaging reader (Biotek).

**Intracellular lipid staining.**
For fluorescence microscopy staining of intracellular lipids, cells were seeded and cultured during 48 h before staining. Cells were fixed 15 min at RT with a 4% formaldehyde solution, washed twice with water before a 5 min incubation with isopropanol 60%. Isopropanol was then removed and Oil-Red-O solution (Millipore Sigma-Aldrich) added on cells for 15 min at RT. Cells were then extensively washed with water to remove the exceeding dye before nucleus counterstaining with NucBlue Fixed Cell Stain ReadyProbes reagent (ThermoFisher Scientific) and observation with a Nikon Eclipse Ts2R microscope (x60). For the quantification of intracellular lipid droplets by flow-cytometry, cells were stained with the BODIPY® 493/503 dye (Tocris Bio-Techne) after 48 h of culture. The cells were washed with PBS before being incubated for 5 min with a 5 µM BODIPY solution in PBS at 37°C. Cells were then washed with PBS before trypsination and FACS analysis. A 7-AAD (BioLegend) staining of dead cells, prior to FACS analysis, allowed gating on living cells.

**Western blot analysis**
Cell lysates from $10^6$ cells were prepared in lysis buffer (1% Triton X-100, 5 mM EDTA in PBS with 1% protease inhibitor cocktail (P8340; Millipore Sigma-Aldrich) and 2 mM orthovanadate). After elimination of insoluble material, proteins were quantified, separated by SDS-PAGE and analyzed by western-blot on PVDF membrane. After saturation of the PVDF membrane in PBS-0.1% Tween 20 supplemented with 5% (w/v) non-fat milk powder, blots were incubated 1 h at room temperature with primary antibody in PBS-0.1% Tween 20 (1:2,000 dilution). Incubation with secondary antibody was performed after washing for 1 h at room temperature. HRP-labelled anti-goat (Santa Cruz Biotechnology), anti-rabbit (A0545, Millipore Sigma-Aldrich) or anti-mouse (Jackson ImmunoResearch Laboratories) antibodies were diluted 20,000 folds and detected by enhanced chemiluminescence reagents according to the manufacturer's instructions (SuperSignal Chemiluminescent Substrate, Thermo Fisher Scientific). Primary antibodies used for immunoblotting included mouse monoclonal antibody against human GCK (clone G-6, Santa Cruz Biotechnology), rabbit monoclonal antibody against human HK2 (Clone C64G5, Cell Signaling Technology), rabbit monoclonal antibody against human HK1 (C35C4, Cell Signaling), rabbit polyclonal antibody against human HK3 (HPA056743, Millipore Sigma-Aldrich), goat polyclonal antibody against human ACLY (SAB2500845, Millipore Sigma-Aldrich), rabbit polyclonal antibody against human pACLY (phospho S455, Cell Signaling Technology), rabbit monoclonal antibody against human PDH (C54G1, Cell Signaling Technology), rabbit monoclonal antibody against human pPDH E1-alpha subunit (phospho S293, Abcam) and goat polyclonal antibody against human PC (SAB2500845, Millipore Sigma-Aldrich).

**Human NK cell purification**
PBMCs were isolated by standard density gradient centrifugation on Ficoll-Hypaque (Eurobio). Mononuclear cells were separated from peripheral blood lymphocytes (PBLs) by centrifugation on a 50% Percoll solution (GE Healthcare). NK cells were purified from PBLs by immunomagnetic depletion using pan-mouse IgG Dynabeads (Thermo Fisher Scientific) with a cocktail of depleting monoclonal antibodies: anti-CD19 (4G7 hybridoma), anti-CD3 (OKT3 hybridoma, ATCC, Manassas, VA, USA), anti-CD4, anti-CD14 and anti-glycophorin A (all from Beckman Coulter). NK purity was >70% as assessed by CD56 labelling (data not shown).

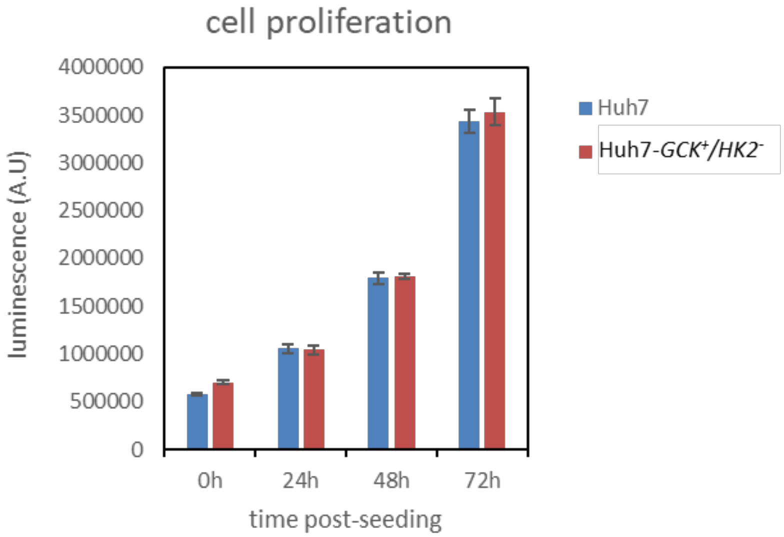


**Supplementary Figure 1.** Proliferation of Huh7 and Huh7-*GCK*$^{+}$/*HK2*$^{-}$. Cells were seeded in 24 well-plate under standard growth conditions and cellular proliferation was determined at time 0, 24, 48 and 72h post-seeding using the CellTiter-Glo® Luminescent Cell Viability Assay (Promega). Luminescence were quantified with an Infinite M200 microplate reader (TECAN). Means ± SEM are presented (n=3).

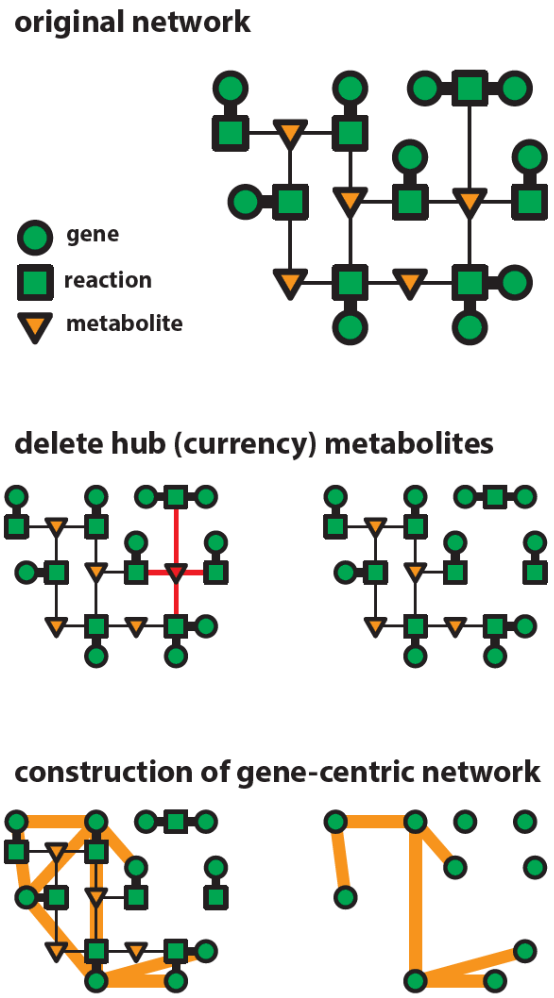


**Supplementary Figure 2.** The original network was extracted from a genome-scale metabolic model where metabolites are connected by reactions that are catalyzed by enzymes. The high-degree nodes ('currency metabolites' like H2O, ATP, etc.) which are not informative about the network organization of the metabolic systems were eliminated before interpreting the network architecture. The removal of different percentages of currency metabolites was tested to obtain reaction-centric graphs. Then gene-reaction associations were used to generate a gene-centric metabolic network with the highest connectivity (see Fig. 2e).

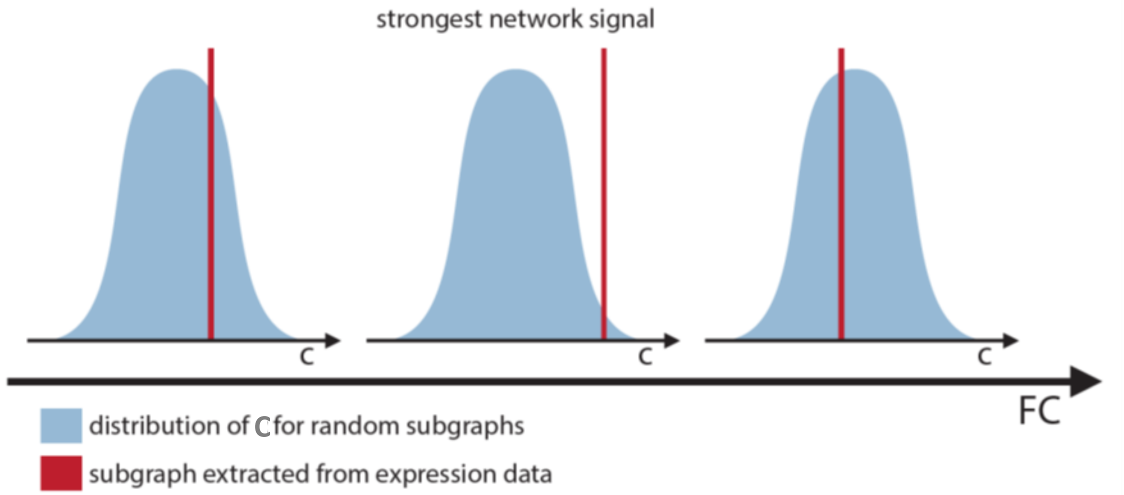


**Supplementary Figure 3.** Given a set S of differentially expressed genes and the gene-centric metabolic network G, we analyzed the subgraph of G spanned by all genes in S. The average clustering coefficient C of connected nodes in these subgraphs serves as a measure of the connectivity of this subgraph (see Supplementary Figure 4). Different FC-expression thresholds of S were applied to calculate C. The FC-expression threshold providing the highest difference of C in comparison to clustering coefficient obtained for a random subgraph (i.e. giving the strongest network signal) was set to calculate the metabolic network coherence MC presented in Supplementary Figure 4).

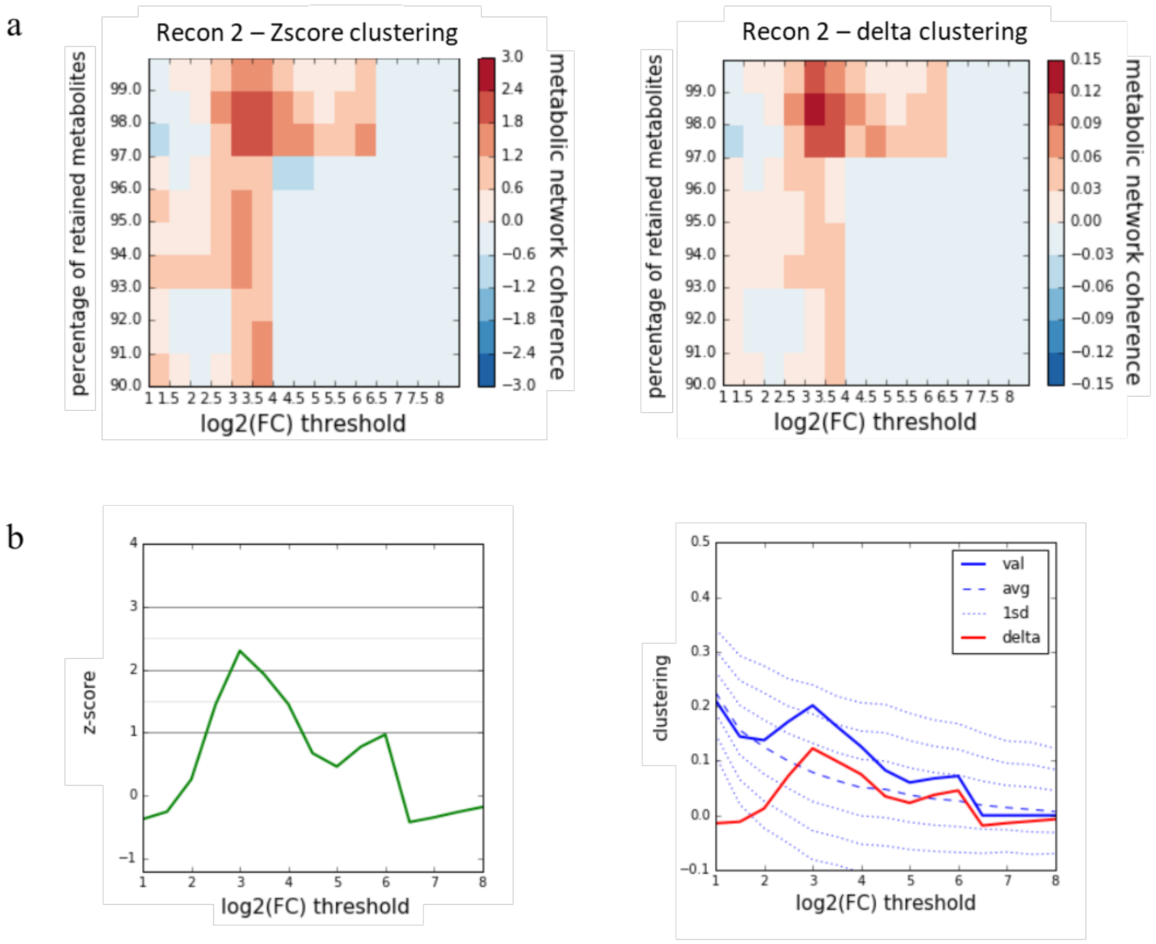


**Supplementary Figure 4. a** Heatmap of Z-scores (left panel) and Delta-scores (right panel) for clustering coefficients obtained when mapping differentially expressed genes to human metabolic model Recon2. Values range from highest in red to lowest in blue as a function of the percentage of retained metabolites (after removal of currency metabolites) and the Fold-Change threshold. **b** Z-score (left panel) and clustering coefficient (right panel) obtained for a currency of 98% (2% of the most connected metabolites removed). Highest Z-score and clustering coefficient C are obtained at log2(FC) threshold >3. This optimal fold-change threshold was used to construct the network presented in Fig. 2e.

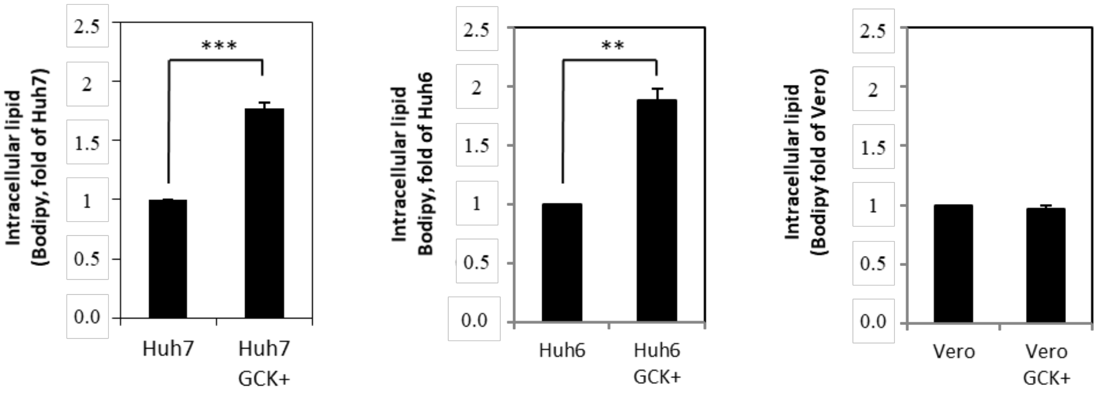


**Supplementary Figure 5.** Intracellular lipid content of two hepatic and one renal cell lines was analyzed after stable re-expression of *GCK*. Parental Huh6, Huh7 or Vero cells were transduced with lentiviruses for *GCK* expression using a pLEX-*GCK* construct as described in material and methods. Cells were then cultured for 7 days in the presence of puromycin to select transduced cells before amplification. Cells were stained for their intracellular lipid content using BODIPY 493/503 dye and analyzed by flow-cytometry. Means ± SEM of fluorescence normalized to the corresponding parental cell line are presented.

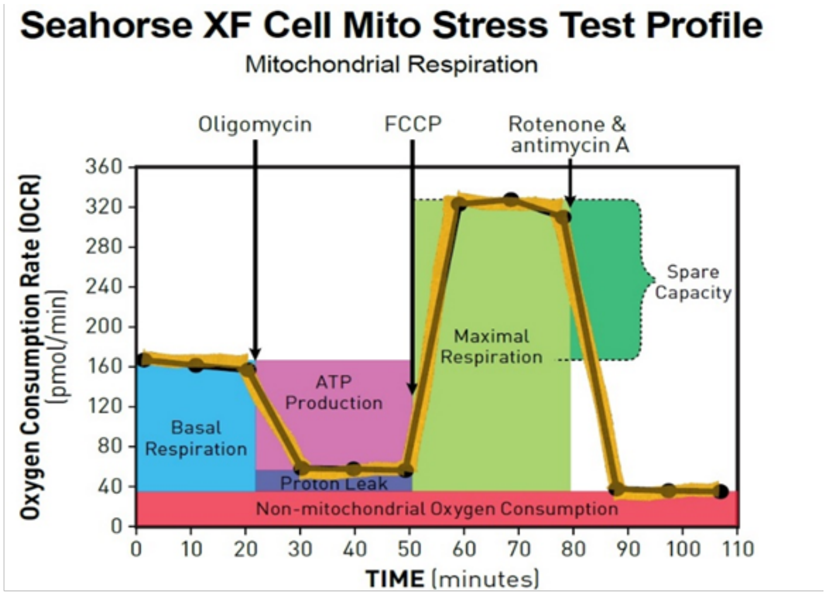


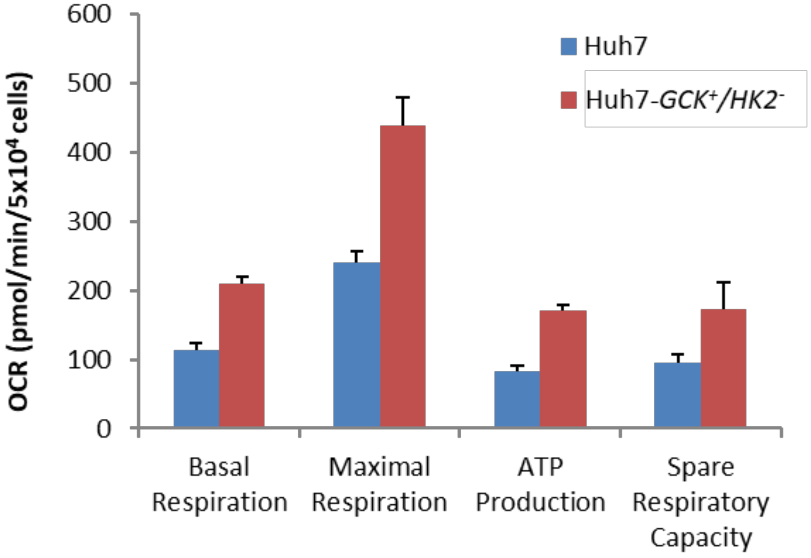


**Supplementary Figure 6.** Basal respiration, Maximal Respiration, ATP production and Spare Respiration Capacity calculated from OCR data generated with Seahorse analyzer (cf. Fig. 5n), as preconized in Seahorse XF cell Mito Stress Test. The results are presented as means ± SEM (n=5).

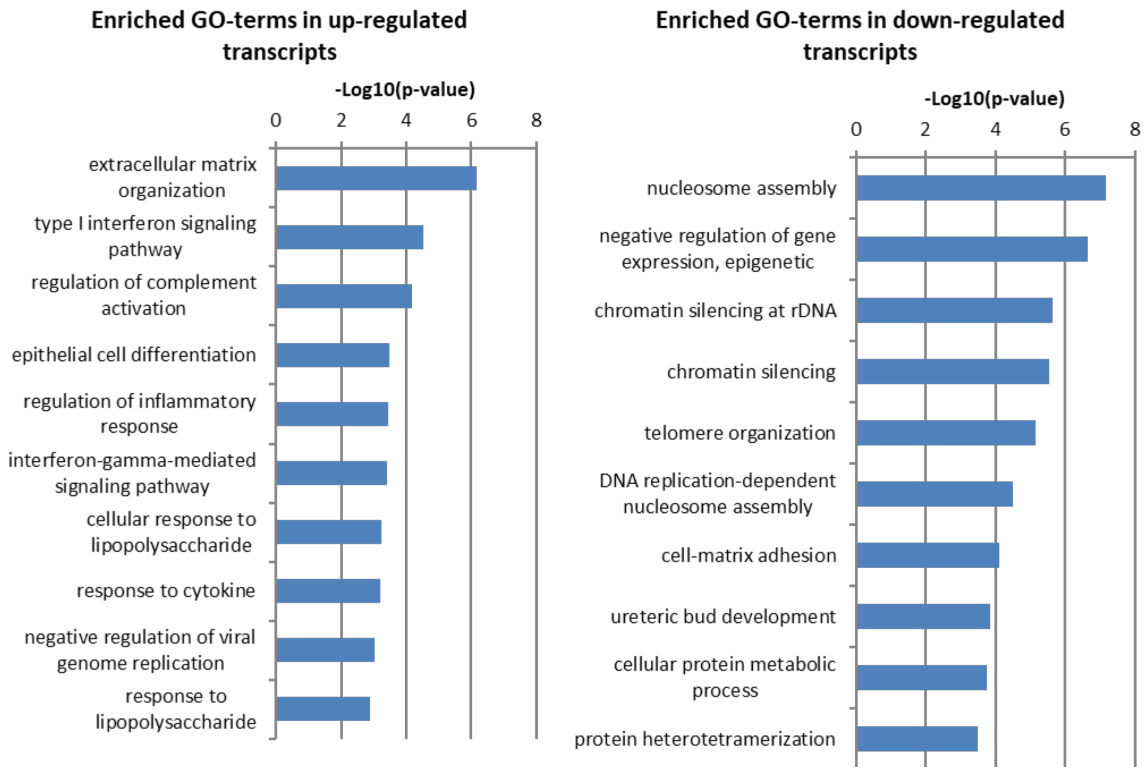


**Supplementary Figure 7.** GO enrichment analysis of transcripts significantly up- and down-regulated in Huh7-*GCK*$^{+}$/*HK2*$^{-}$ vs Huh7 cells determined by DESeq2 analysis of transcriptomic data. Transcripts lists were submitted to the Functional Annotation Tool of the online knowledge base DAVID Bioinformatics Resources 6.8, NIAID/NIH. In DAVID, Fisher's Exact test is adopted to measure the gene-enrichment in annotation terms. Parameters were adjusted to a minimal count belonging to the annotation of 10 and EASE score (one-tail Fisher Exact Probability Value) threshold of 0.01.

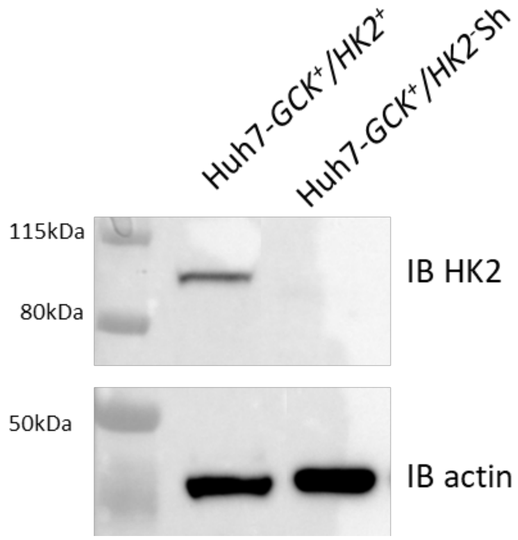


**Supplementary Figure 8.** Western Blot analysis of HK-II protein expression in Huh7 cells transduced for *GCK* expression before (Huh7-*GCK*$^{+}$/*HK2*$^{+}$) and after extinction by ShRNA (Huh7-*GCK*$^{+}$/*HK2*$^{-}$Sh). Immunoblot of actin was performed on the same blot for normalization.

a

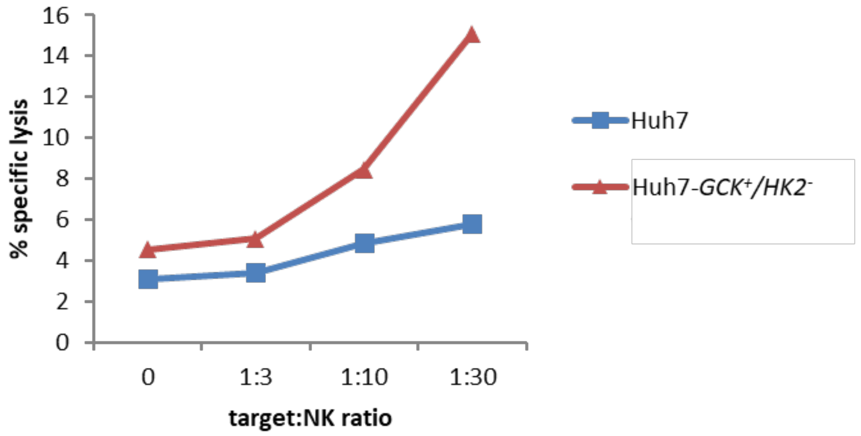


b

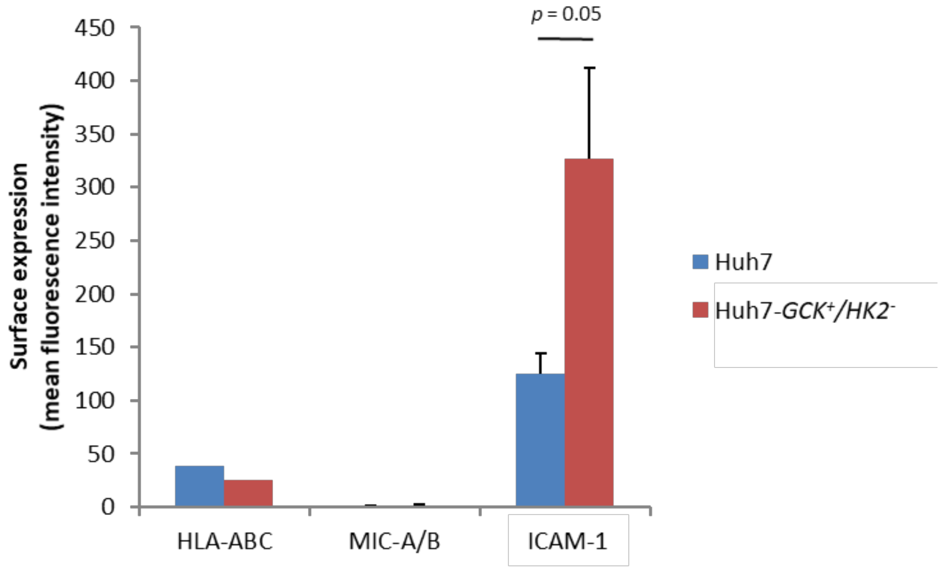


**Supplementary Figure 9. a** NK cell mediated lysis of Huh7 or Huh7-$GCK^{+}/HK2^{-}$ . Hepatoma cells were seeded 24 h before addition of IL2-preactivated NK cells for 4 h at effector to target (E:T) ratio of 0, 3, 10 or 30. After harvesting, cells were stained with propidium iodide (PI) and analyzed by flow cytometry. Lysis was determined by the percentage of $PI^{+}$ cells on gated hepatocytes. **b** Surface expression of HLA-A,B,C, MIC-A/B and ICAM-1 on Huh7 and Huh7-$GCK^{+}/HK2^{-}$ cells analyzed by Flow cytometry after cell surface immunolabelling with anti-HLA-ABC-PE (BD Biosciences), anti-MIC-A/B-PE (BD Biosciences) or anti-ICAM-1-FITC (Beckman Coulter).

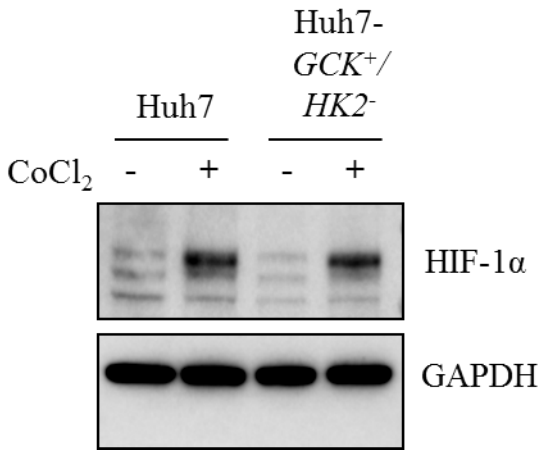


**Supplementary Figure 10**. Western Blot analysis of HIF-1α in Huh7 and Huh7-*GCK*$^{+}$*/HK2*$^{-}$ cells with or without 150 μM $CoCl_2$ treatment to mimic hypoxic conditions. Immunoblot of GAPDH was performed on the same blot for normalization.

**Supplementary Table 1 (separate file).** *HK1, HK2, HK3* and *GCK* expression levels across HCC biopsies from 365 patients (Data from The Cancer Genome Atlas - TCGA) used to construct Figure 1.

**Supplementary Table 2 (separate file).** Transcriptomes of Huh7 and Huh7-$GCK^{+}/HK2^{-}$ cells. Genes expression of the two cell lines was analyzed by Next-Generation Sequencing (NGS). Reads were mapped on the reference genome Homo sapiens GRCh37/hg19 and analyzed using DESeq2 method. Differential expression of genes between Huh7 and Huh7-$GCK^{+}/HK2^{-}$ cell lines were used in figures 2d,e, 3a-c, 5j and 6a,b. Entire Raw data are available in Gene Expression Omnibus database with the accession number GSE144214.

**Supplementary Table 3 (separate file).** The complete list of genes included in top 5 enriched functions identified by Ingenuity Pathway Analysis (IPA). Enrichment of molecular and cellular functions in differentially expressed genes between Huh7 and Huh7-$GCK^{+}/HK2^{-}$ cell lines. The list of transcripts differentially expressed in Huh7 and Huh7-$GCK^{+}/HK2^{-}$ cell lines was analyzed by gene set enrichment analysis IPA (Build version: 486617M, Qiagen) weighted by their corresponding fold change and *p* value.
